# Fast Pangenome Matching Statistics in Compressed Space with Move Structures

**DOI:** 10.64898/2026.11.10.765781

**Authors:** Nathaniel K. Brown, Ahsan Sanaullah, Pramesh Shakya, Shaojie Zhang, Ben Langmead

## Abstract

Matching statistics and supermaximal exact matches (SMEMs) are key inputs for sequence homology search and classification. Methods such as MONI, PHONI, and ropebwt3 compute these statistics using compressed pangenome indexes, but are substantially slower than methods that compute approximations of matching statistics, such as pseudo-matching lengths (PMLs). PMLs can be computed quickly in a single pass, but SMEM computation can require overlapping forward or backward passes over the query. We present ms2, a suite of algorithms that use “move structures” to compute matching statistics and SMEMs efficiently in compressed space.

Our four algorithms compose move structures for the LF, *Ψ*, *ϕ*, and *ϕ^−^*^1^ permutations with additional longest common prefix (LCP) array summaries and samples. The algorithms offer different tradeoffs among speed, index size, and support for reporting SMEM offsets. They inherit the move structure’s fast, constant-time operations and achieve high throughput by leveraging the move structure’s locality of reference. Thanks to recent advances, move structures can be constructed in optimal time and compressed space, including the LCP information needed for “match recovery” during SMEM computation.

On 100 haplotypes of human chromosome 22, ms2’s fastest algorithm computes matching statistics 35 to 48 times as fast as PHONI’s online approach, while using an index of similar size. On the 472-haplotype HPRC2 pangenome and human reads, three ms2 modes achieve speedups of at least 22, 19, and 15 times over ropebwt3, whose index is about 15% the size of ms2’s fastest mode. When additionally reporting SMEMs and their offsets, ms2 achieves speedups of 25 to 34 times over ropebwt3. Together, these advances enable online computation of matching statistics and SMEMs at a throughput that supports real-time processing of approximately 21 PromethION nanopore flow cells on a single thread.

The ms2 software is open source and available at https://github.com/langmead-lab/ms2. It builds on the open-source TeraTools suite (https://github.com/ucfcbb/TeraTools) and Orbit library (https://github.com/drnatebrown/orbit). Experimental scripts and other details are described in: https://github.com/langmead-lab/ms2-experiments.

## 1 Introduction

Matching statistics and supermaximal exact matches (SMEMs) are high-fidelity descriptions of maximal exact sequence matches between a query and a reference. Their lengths provide evidence for metagenomic sequence classification [5, 56, 22, 18], while their reference locations supply seeds for read and genome alignment [54, 38, 45]. Compared to *k*-mer-based methods, SMEM-based methods can be more powerful, since SMEMs give answers to *k*-mer presence/absence questions for any value of *k* [34, 6].

As reference collections grow, conventional indexes grow proportionally to the amount of sequence in the collection. Compressed indexes, on the other hand, exploit repetition among genomes and scale only with the non-redundant sequence content, enabling indexes that fit in memory even for large pangenomes [30, 38]. Still, recovering the lengths of maximal exact matches has remained challenging in compressed space.

While MONI and PHONI are capable of computing matching statistics, they do this with relatively heavy auxiliary structures. MONI uses a random-access data structure built over the pangenome, which it queries in a separate second pass [49]. PHONI augments grammar-supported text access with longest common extension (LCE) support, allowing it to compute matching statistics in an online fashion [12]. These randomaccess and LCE operations remain a bottleneck despite subsequent optimizations [42, 31]. Bidirectional approaches like ropebwt3 avoid those particular auxiliary structures, but expend computational effort repeatedly extending overlapping matches, potentially requiring a quadratic-in-query-length number of steps [38, 14]. SPUMONI reduces time and space by computing pseudo-matching lengths (PMLs) without random text access [5, 4], but that method computes neither full matching statistics nor SMEMs.

Nishimoto and Tabei’s *move structure* [43] has been used extensively to accelerate *Burrows-Wheeler transform* (BWT [19]) based compressed text indexing [15, 11, 24, 35] and pangenome queries [51, 48, 22, 18]. In particular, Movi uses move structures to accelerate PML computation through constant-time navigation and unique latency-hiding optimizations [56, 57]. To date, no one has applied the move structure to the problems of computing matching statistics and SMEMs in a fast, online fashion. The challenge is to approach the throughput of PML computation while recovering exact matching statistics and SMEMs online.

We present ms2, for *move structure matching statistics*, a suite of four algorithms that recover exact match lengths directly from the compressed index. All four compute matching statistics and SMEMs in a single right-to-left pass, recovering the matching-statistics lengths through the use of sampled longest common prefix array (LCP) minima, local text extraction, or LCP enumeration. The fourth strategy is a hybrid that interleaves computation from the extraction and enumeration strategies, finishing as soon as either resolves the crucial “recovery” step. These methods give explicit recovery bounds and achieve highly advantageous tradeoffs among query time, index space, and support for reporting match occurrences.

ms2’s speed and flexibility come from its use of move structure indexes, which we compose in novel ways to achieve flexible and fast navigation in both text coordinates (*ϕ* and *ϕ^−^*^1^) and lexicographical coordinates (LF and *Ψ*). The Orbit library [16, 17] enables these compositions through shared storage, integrated auxiliary data, and economical allocation of bits to each field of the structure. Orbit reduces storage while promoting locality of reference, which in turn allows ms2 to benefit from aggressive latency-hiding approaches. Building ms2 indexes is also quite efficient, since TeraTools [50] and Orbit provide optimal-time construction of the underlying move structures and LCP samples in compressed space.

We evaluate ms2’s efficiency in various analysis scenarios, including with very large pangenomes. We find that the sampled-minima method (ms2-min) approaches the speed of an ideal “oracle” baseline where all recovery steps are simply recalled from a precomputed list. On 100 chromosome 22 haplotypes, the ms2 methods compute matching statistics 35 to 48 times faster than PHONI with similar memory. On the 472-haplotype HPRC2 pangenome, ms2 is at least 22 times faster than ropebwt3 on human reads, though ropebwt3’s index is about 15% as large. When reporting SMEM occurrences, ms2 is 8.8 to 11 times faster than ropebwt3 when they are configured to use a similar memory footprint. The single-thread throughput of ms2’s fastest mode on nanopore reads can keep up with approximately 21 PromethION flow cells.

## 2 Preliminaries

### 2.1 Notation and BWT Indexing

Let *T* [0*..n* − 1] be a text terminated by a unique smallest symbol $, with other characters drawn from an ordered alphabet *Σ* of constant size |*Σ*| = *σ*. Let *P* [0*..m* − 1] be a pattern over *Σ*, and *lcp*(*u, v*) denote the longest common prefix length of strings *u* and *v*. We assume the word-RAM model with machine words of size *w* = *Θ*(log *n*) and measure space in words unless otherwise specified.

The suffix array SA[0*..n* − 1] stores the starting positions of *T*’s suffixes in lexicographic order. The *Burrows-Wheeler Transform* (BWT) [19] is a permutation of *T* such that BWT[*i*] = *T* [(SA[*i*]−1) mod *n*]. It is navigated backward by the *last-to-first* (LF) mapping, satisfying SA[LF(*i*)] = (SA[*i*] − 1) mod *n*, and forward by its inverse, the *Ψ* (or FL) mapping, satisfying SA[*Ψ* (*i*)] = (SA[*i*] + 1) mod *n*. The *run-length encoded BWT* (RLBWT) compresses the BWT into its *r* maximal equal character runs. The permutation *ϕ* is defined such that *ϕ*(SA[*i*]) = SA[(*i* − 1) mod *n*], with its inverse symmetrically defined as *ϕ^−^*^1^(SA[*i*]) = SA[(*i* + 1) mod *n*]. The *r*-index [30] supports queries in *O*(*r*)-space by augmenting the RLBWT with compressed representations of LF and *ϕ*, alongside SA samples at BWT run boundaries.

The *longest common prefix array* (LCP) stores the *lcp* between lexicographically adjacent suffixes, with LCP[0] = 0 and LCP[*i*] = *lcp*(*T* [SA[*i* − 1]*..n* − 1]*, T* [SA[*i*]*..n* − 1]). The *r* values LCP[*i*] where *i* starts a BWT run are the *irreducible* LCPs, from which all remaining LCP values can be derived. For suffix array ranks *p < q*, a *longest common extension* (LCE) query is defined as LCE(SA[*p*], SA[*q*]) = *lcp*(*T* [SA[*p*]*..n* − 1]*, T* [SA[*q*]*..n* − 1]), which can be evaluated via a range minimum query (RMQ) over the LCP array as min*_p<k≤q_* LCP[*k*]. *Previous* and *next smaller value* queries are defined as PSV(*i*) = max{*k < i* | LCP[*k*] *<* LCP[*i*]} and NSV(*i*) = min{*k > i* | LCP[*k*] *<* LCP[*i*]}.

### 2.2 Move Structures

Nishimoto and Tabei’s *move structure* [43] represents a permutation *π* using *O*(*b*)-space, where *b* counts the number of maximal intervals on which *π*(*i* + 1) = *π*(*i*) + 1. For a move structure M*_π_*, a *move query* M*_π_.*move(*i, j*) = (*π*(*i*)*, j^′^*) maps position *i* in interval *j* to *π*(*i*) in interval *j^′^*. Balancing selectively splits these intervals to support move queries in *O*(1)-time while retaining *O*(*b*)-space. The permutations LF, *Ψ*, *ϕ*, and *ϕ^−^*^1^ each decompose into *r* such intervals, yielding constant-time move queries in *O*(*r*)-space [43]. In particular, maximal LF intervals correspond directly to BWT runs. The move structure’s strong locality of reference also enables cache-efficient queries and latency hiding in practice, as demonstrated by Movi [56, 57]. Length capping splits intervals further to bound their lengths by *O*(*n/r*) without changing these guarantees [16]. We call the resulting intervals *move intervals*.

Augmenting *ϕ* with irreducible LCP samples supports enumeration of adjacent SA and LCP values in *O*(1)-time per step using the move structure [30, 53, 51]. Sanaullah et al.’s TeraTools [50] computes these samples from the RLBWT in *O*(*n* + *r* log *r*)-time and *O*(*r*)-space, with the *r* log *r* term arising from move structure balancing. Brown and Langmead’s [16] length capping removes the need for balancing during full permutation traversals, while Brown et al. [17] subsequently give an *O*(*r*)-time balancing algorithm. Together, these results construct balanced LF*/Ψ* structures in *O*(*r*)-time, and balanced *ϕ/ϕ^−^*^1^ structures and irreducible LCP samples in *O*(*n*)-time, from the RLBWT, all in *O*(*r*)-space.

### 2.3 Matching Statistics and SMEMs

The *matching statistics* (MSs) of a pattern *P* [0*..m* − 1] with respect to a text *T* [0*..n* − 1] are defined as an array MS[0*..m* − 1] where MS[*i*] is the maximum length *ℓ* such that *P* [*i..i* + *ℓ* − 1] occurs in *T*. Some definitions of MSs also require a text position; when required, Pos[*i*] stores the starting position of an arbitrary occurrence of *P* [*i..i* + MS[*i*] − 1] in *T*.

A *supermaximal exact match* (SMEM) of *P* with respect to *T* is a substring *P* [*i..j*] that occurs in *T* but does not occur when extended in either direction. Formally, if *i >* 0, then *P* [*i* − 1*..j*] does not occur in *T*, and if *j < m* − 1, then *P* [*i..j* + 1] does not occur in *T*. Let *ℓ_i_* = MS[*i*]. Then *P* [*i..i* + *ℓ_i_* − 1] is an SMEM if and only if *ℓ_i_ >* 0 and either *i* = 0 or *ℓ_i−_*_1_ ≤ *ℓ_i_*. Consequently, any algorithm for computing MSs can report all SMEMs, either during computation or with an additional *O*(*m*)-time scan. We refer to a matching algorithm as *online* if it processes query characters in scan order and reports results incrementally, without access to unread characters.

#### Interval-Based Methods

Classical algorithms compute matching statistics by navigating suffix tree loci [20] or their corresponding suffix array intervals [2], exploiting suffix links to achieve *O*(*m*)-time with *O*(*n*)-space indexes. Ohlebusch et al. [44] simulate these suffix links in compressed full-text indexes using PSV/ NSV queries over the LCP array. However, these auxiliary navigation structures do not directly permit *O*(*r*)-space representations, though Gagie et al. [30] can answer PSV/NSV and RMQ queries in *O*(*r* log(*n/r*))-space with logarithmic query times.

Li’s [37] forward-backward approach sidesteps these navigational queries by finding SMEMs directly through repeated extensions in both directions. RLBWT-based bidirectional indexes support these extensions in *O*(*r* + *r*)-space [7, 24, 26], where *r* counts the BWT runs of the reversed text. Li’s ropebwt3 [38] uses this strategy to find SMEMs and count their occurrences, and additionally supports locating selected matches. However, forward-backward search performs *Θ*(*m*^2^) extensions in the worst case [14]. Boyer-Moore-Li skipping [27], implemented in ropebwt3, and *k*-mer filtering with KeBaB [14] reduce work when seeking only long SMEMs, but do not completely eliminate the repeated traversal of long, overlapping matches.

#### Single-Candidate Methods

Bannai et al. [8] augment the *r*-index with *thresholds* to maintain one suffix array candidate rather than a full interval. A right-to-left pass uses an *O*(*r*)-space index to extend the candidate via LF when the next character matches. On mismatch, precomputed thresholds select between the nearest preceding and following occurrences of the required character in the BWT, *repositioning* to the candidate which retains the longest match. A subsequent left-to-right pass recovers match lengths using random access to the text. Rossi et al.’s MONI [49] implements this approach, constructing thresholds efficiently through prefix-free parsing [13] and supporting text access with a grammar compressed text representation. With a grammar of size *g* supporting logarithmic-time text access, this approach computes MSs in *O*(*m* log *n*)-time using an *O*(*r* + *g*)-space index.

To achieve an online algorithm, PHONI [12] uses grammar-supported LCE queries to select candidates and recover exact lengths in a single pass. However, these LCE queries dominate query time in practice [42, 31]. Precomputed LCE bounds as used in AugPHONI [42] and delayed *lcp* evaluation [31] reduce this cost, but text access remains a bottleneck. To reduce query time and index space, SPUMONI [5, 4] omits the grammar and suffix array samples while retaining repositioning thresholds, resetting the length on mismatch to compute *pseudo-matching lengths* (PMLs). Movi implements this with move structures; its constant-time configuration supports PML computation in *O*(*m*)-time with an *O*(*r*)-space index [56].

## 3 Methods

Our ms2 algorithms adopt the single-candidate framework of Bannai et al. [8], scanning the pattern from right to left. We distinguish two operations: *match extension*, which extends the current candidate to the left by one character, and *match recovery*, which selects a replacement candidate that permits extension while retaining as much of the current match as possible. Our algorithms additionally determine the retained length during recovery, enabling online computation of exact matching statistics and SMEMs in the same right-to-left pass. They share the same extension mechanism but implement match recovery differently.

### Match Extension

At each step, we maintain a suffix array rank *p* and match length *ℓ* = MS[*i*], taking MS[*m*] = 0, with

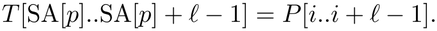

Initially, *i* = *m*, *ℓ* = 0, and *p* is arbitrary. If BWT[*p*] = *P* [*i* − 1], we perform *match extension*: set *p* = LF(*p*), increment *ℓ*, and set MS[*i* − 1] = *ℓ*.

We implement extension with a balanced LF move structure using *O*(*r*)-space. Its move intervals subdivide BWT runs, and we store the corresponding run character with each interval to support BWT[*p*] access. The move query M_LF_.move(*p, j*) = (LF(*p*)*, j^′^*) updates both the candidate rank and its containing interval in *O*(1)-time. Thus, match extension steps take *O*(1)-time each and *O*(*m*)-time in total; however, there can be *Θ*(*m*) mismatch cases, so each algorithm’s total query complexity depends on its match recovery cost.

### Match Recovery

When BWT[*p*] = *P* [*i* − 1], we perform *match recovery*: select a replacement candidate that permits extension while retaining the longest possible prefix of the current match. By lexicographic order, it suffices to consider *repositioning* to the nearest preceding and following occurrences of *P* [*i* − 1] in the BWT. We locate these in *O*(1)-time using per-character links between move intervals in *O*(*rσ*) = *O*(*r*)-space. We compute the retained lengths at these candidates to select a replacement and update the match length.

For an eligible candidate of rank *q < p*, the retained length is an LCE capped by the current match length:

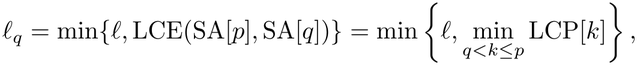

with *q > p* handled symmetrically. We select a candidate maximizing *ℓ_q_*, then extend by setting *p* = LF(*q*) and *ℓ* = *ℓ_q_* + 1, yielding MS[*i* − 1] = *ℓ*. If either candidate retains all *ℓ* characters, we select it without further evaluating the other candidate (*full-match retention*), breaking ties arbitrarily. If *P* [*i* − 1] does not occur in *T*, we instead set *ℓ* = MS[*i* − 1] = 0. After either match extension or recovery, we decrement *i* and continue.

Candidate lookup and LF extension take *O*(1)-time, leaving the capped LCE queries as the main recovery cost. We next present four methods for answering these queries by composing move structures with auxiliary data, without a separate random-access representation of the text. The descriptions below establish the theoretical bounds; practical modifications are detailed under *Implementation*.

### 3.1 Match Recovery Methods

#### LCP Minima Sampling

Our first method, ms2-min, samples the LCP minima needed for match recovery, reporting matching statistics and SMEMs, but not text offsets. Index the move intervals in order, and let interval *j* span [*s_j_, e_j_*], with boundary value *b_j_* = LCP[*s_j_*] and interior minimum *µ_j_* = min*_sj_ <k≤e_j_* LCP[*k*]. Since replacement candidates lie at BWT run boundaries, each required LCE range decomposes into a partial interior of the current move interval and a sequence of intervening boundary values and interval interiors (Figure 1). We store the *b_j_* and *µ_j_* values in an *O*(*r*)-space RMQ structure [10, 25], which returns the minimum *R* over the intervening boundary values and interval interiors in *O*(1)-time. During recovery from interval *j*, let *C* = min{*ℓ, R*}. If *C* ≤ *µ_j_*, then *C* is the capped LCE and we omit the directional query (*summary skipping*); otherwise, we compute the directional minimum from *p* within interval *j*.

**Fig. 1.**
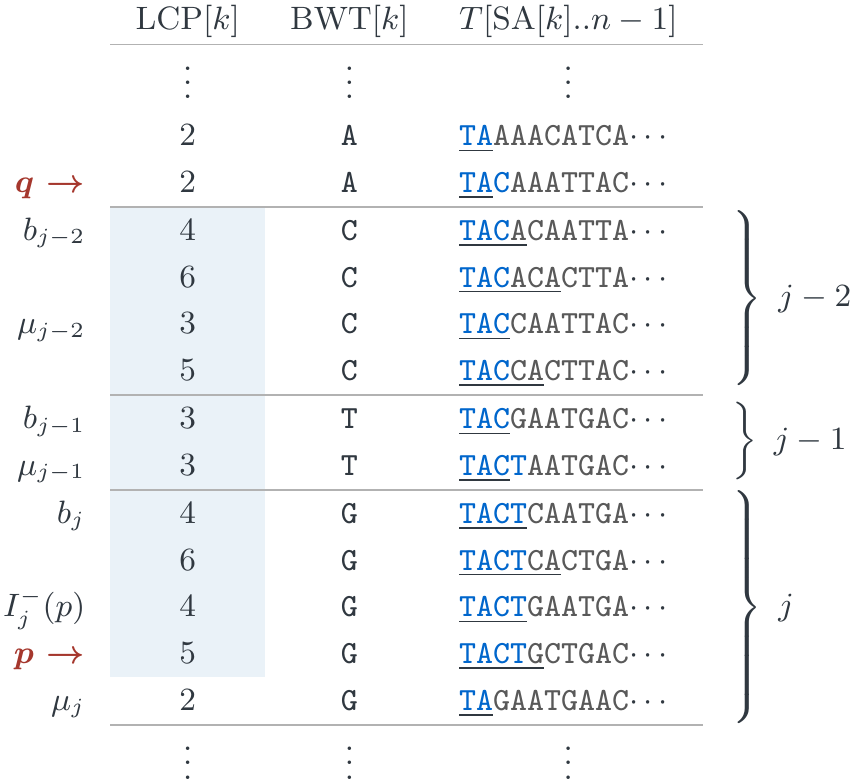
Match recovery from *p* in interval *j*, matching TACT, to the preceding candidate *q* with BWT[*q*] = A. Shading marks the required LCE range LCP[*q* + 1*..p*]; blue prefixes match `TACT`; and underlines span LCP[*k*] characters. Left labels show boundary values, interior minima, and 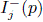; braces identify move intervals. The intervening summaries yield *R* = 3, and *ℓ* = 4 gives *C* = 3. Since *µj* = 2 *< C*, a directional query is required but returns 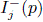 = 4, so recovery retains three characters.

For each interval *j*, we scan LCP[*s_j_* + 1*..e_j_*] in both directions, recording a position and value whenever the running minimum decreases. These samples form an NSV chain from the left and a PSV chain from the right, representing 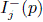 = min*_s <k≤p_* LCP[*k*] and 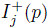 = min*_p<k≤e_* LCP[*k*].

Because each LCE range includes the adjacent boundary, we can safely omit left-chain samples with values at least *b_j_* and right-chain samples with values at least *b_j_*_+1_, possibly eliminating a chain entirely. When a directional query remains, binary search over the *κ* retained samples returns min{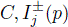} in *O*(log *κ*)-time. Length capping gives *κ* = *O*(*n/r*), yielding *O*(log(*n/r*)) time per recovery and *O*(*m* log(*n/r*)) total query time.

Differential-LCP representations exploit the identity LCP[LF(*i*)] = LCP[*i*] + 1 whenever BWT[*i* − 1] = BWT[*i*] to compress LCP information on repetitive texts [1, 30]. Similarly, expressing sample positions relative to *s_j_* and LCP values relative to *µ_j_* permits sharing repeated or overlapping directional chains. A shared contiguous array with per-direction offsets and lengths supports *O*(1)-time sample access. If this array occupies *S* words, the index uses *O*(*r* + *S*) space and retains the *O*(log(*n/r*)) recovery bound. Given an RLBWT, we construct the boundary values, interior minima, and uncompressed chains in *O*(*n*)-time and *O*(*r*) working space in addition to the output [50, 17]. Our implementation uses interval-relative positions but unnormalized LCP values, merging both directions into variable-length encodings while sharing identical records (Appendix Note A4).

Caching reduces recovery cost when *P* is similar to *T*; formally, if the BWT of their concatenation does not decrease in average run length compared to either string alone, caching directional minima yields *O*(*m*) query time in *O*(*r* + *S* + *m*)-space (Appendix Note A1). Further, a variant of ms2-min can maintain the full SA interval for occurrence counting, taking *O*(log *r* + log(*n/r*)) time per recovery without changing the index space bound and match extension time (Appendix Note A2).

#### Text Extraction with Ψ

Our second method, ms2-psi, computes the capped LCE directly by extracting and comparing characters. We compose the LF structure with a *Ψ* move structure. Given a candidate rank *q* and its LF interval *j*, stored links identify *q*’s containing *Ψ* interval *h* in *O*(1)-time. Starting at *k* = *q*, cumulative character counts identify *T* [SA[*k*]], and queries M*_Ψ_.*move(*k, h*) = (*Ψ* (*k*)*, h^′^*) advance to the next character. We compare these characters with *P* [*i..i*+*ℓ* − 1], stopping at the first mismatch or after *ℓ* characters to obtain *ℓ_q_* (*extraction stopping*). At *O*(1)-time per character, each match recovery takes *O*(*ℓ_q_*) time.

The total query time is *O*(*m* + *_i_* MS[*i*]), or *O*(*m*^2^) in the worst case. Short matches limit the forward comparisons, making this approach favorable for patterns with low matching statistics. The two move structures and their navigation support use *O*(*r*)-space. Extraction revisits processed pattern characters, requiring up to *O*(*m*) characters of query buffering in addition to the index. By default, ms2-psi does not report text offsets, but adding SA samples at BWT run boundaries provides SA[*q*] for each replacement candidate. We maintain the text position by decrementing it after each LF match extension, as described by Bannai et al. [8], and report Pos[*i* − 1] alongside MS[*i* − 1].

#### LCP Enumeration with ϕ

Our third method, ms2-phi, uses the same interval summary framework as ms2-min, including the boundary values *b_j_*, interior minima *µ_j_*, RMQ support, and summary skipping condition. However, it replaces directional sample chains with direct enumeration of adjacent LCP values. We compose the LF structure with a *ϕ* move structure alongside irreducible LCP samples, supporting enumeration in *O*(1)-time per value in *O*(*r*)-space [30, 53, 51]. We maintain *x* = SA[*p*] and its containing *ϕ* interval *h* alongside (*p, j*). Each LF interval head links to a *ϕ* interval starting at its corresponding SA value [43]. One *ϕ* move from the position linked at the following LF interval head reaches the suffix at the current interval tail. These links update *x* and *h* after selecting a replacement candidate.

When *C > µ_j_*, we compute the required 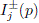 by enumeration. For 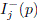, we enumerate ranks from *p* down to *s_j_* +1; for 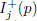, we start at the interval tail *e_j_* and enumerate down to *p* +1. Each move M*_ϕ_.*move(*x, h*) = (*ϕ*(*x*)*, h^′^*) yields an LCP value by subtracting *x*’s interval offset from its irreducible sample [29]. We stop when the running minimum reaches *µ_j_* or the required range is exhausted (*enumeration stopping*). Each LF extension decrements *x* modulo *n* and updates *h* only when crossing an interval boundary [50].

Length capping bounds each enumeration by *O*(*n/r*), yielding *O*(*m* · *n/r*) query time and *O*(*r*)-space. The caching and interval-maintenance variants of ms2-min also extend to ms2-phi. Under the same pattern similarity condition, caching the enumerated directional minima yields *O*(*m*) query time in *O*(*r* + *m*) total space (Appendix Note A1). Maintaining the full SA interval for occurrence counting takes *O*(log *r* + *n/r*) time per recovery without changing the index space bound and match extension time (Appendix Note A2).

The maintained text position supplies Pos[*i*]. We report all occurrences by enumerating SA and LCP values, using MS[*i*] to extend to the correct match interval. The *ϕ* structure enumerates directly from *p* toward smaller ranks; toward larger ranks, traversal starts at interval tails and may scan unmatched portions of length capped intervals, giving *O*(occ + *n/r*) time to report all occ occurrences. Additionally composing a *ϕ^−^*^1^ structure permits direct enumeration in both directions in *O*(occ)-time, while retaining *O*(*r*)-space.

#### Dual Approach

Our fourth method, ms2-dual, combines ms2-psi and ms2-phi. Short matches limit *Ψ* comparisons, whereas short directional enumeration limits *ϕ* steps. For each replacement candidate, we alternate between the two traversals until one completes, incurring at most twice the cost of the faster traversal. This yields *O*(min{MS[*i*]*, n/r*})-time on match recovery and 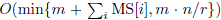 total query time with an *O*(*r*)-space index. As in ms2-psi, extraction requires buffering up to *O*(*m*) pattern characters. Locating support is inherited from ms2-phi. Successful *Ψ* comparisons raise a candidate’s lower bound, while LCP enumeration lowers its upper bound. Comparing these bounds across candidates can discard one before its exact retained length is known (*candidate discarding*); see Appendix Note A5.

### 3.2 Implementation

We use TeraTools to construct RLBWT-derived permutation data and LCP samples [50], and Orbit to construct and represent move structures [16, 17]. Orbit supports user-defined metadata columns packed alongside move intervals, which we use to store characters, navigation links, and LCP samples. Our implementation shares storage between LF and its inverse *Ψ* using the *invertible move structure*, described in Appendix Note A3. Figure 2 shows the components of the four methods.

**Fig. 2.**
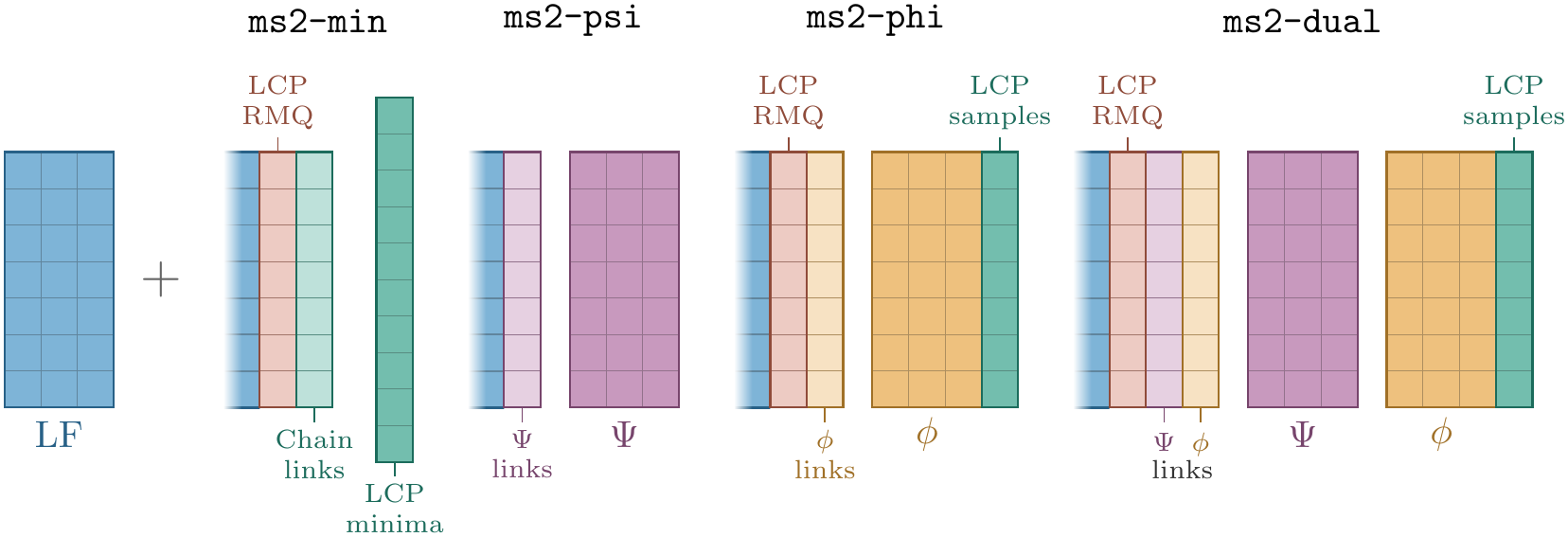
Move structures and supporting data in the theoretical formulations of the four ms2 methods. Dimensions are schematic: field widths and numbers of move intervals may vary. Appendix Table A1 gives the storage layout of each move structure.

We replace the *O*(1)-time character links and RMQ support with linear scans over interval characters and LCP summaries. For ms2-min, we also scan retained samples linearly rather than using binary search. These choices are efficient in practice but do not preserve the preceding query-time guarantees. We interleave independent pattern queries to hide memory latency [56] and parallelize across patterns. ms2-psi, ms2-phi and ms2-dual support occurrence enumeration using *ϕ* and *ϕ^−^*^1^ move structures. Appendix Notes A4 and A5 describe storage layouts and query execution, including the recovery stopping conditions. Our code is available at https://github.com/langmead-lab/ms2.

## 4 Results

### Measurements

All experiments were conducted on Amazon EC2 `i4i.metal` computers. Each computer has 32-core Intel Xeon Platinum 8375C processors and 1 TiB of DRAM over two NUMA nodes. Tools were bound with `numactl` to one NUMA node to avoid NUMA effects. Each measured value is a median from three separate runs. We measured index loading separately from querying and report speed over query time only. Memory is measured as peak resident set size (RSS) and reported in GiB (2^30^ bytes). Appendix Note A6 provides more detail, and the commands, seeds, input checksums and per-run records of every experiment are in the ms2-experiments repository (https://github.com/langmead-lab/ms2-experiments, tag `paper-v1.1`).

### Recovery cost for ms2 modes

We first measured recovery cost in each ms2 mode through query time and move structure operations per query base (Fig. 3). We compared each mode with an oracle that replaces candidate selection and length recovery with results recorded by an earlier run, providing a recovery-free baseline. We used two indexes. First, the “mtb152” index consists of 144 out of the 152 *M. tuberculosis* genomes generated and used previously [41, 21]. We obtained these from a ropebwt3 index distributed by Li [39], with the first 8 genomes held out as potential query sequences. Second, the “HPRC2” index consists of the 464 haplotypes of release 2 of the HPRC [40] together with CHM13 [3], GRCh38 [52], CN1 [55], YAO [33] and KSA001 [36] (472 haplotypes in all). We queried each index with two types of queries: pieces of whole genomes of the same species but absent from the index (“genomes”) and PacBio HiFi reads simulated from those same genomes (“PacBio reads”). For mtb152, the query genomes were the eight held-out genomes and their reverse complements. For HPRC2, the query genome was haplotype 1 of APR008 from the Arab Pangenome Reference (GCA_050494745.1), which is not present in HPRC2. We cut query genomes into 100 kbase pieces and drew a seeded random sample of 600 pieces from each index’s set, with some truncated at sequence ends, leading to 59.1 Mbases of query sequence total for mtb152 and 59.9 Mbases for HPRC2.

**Fig. 3.**
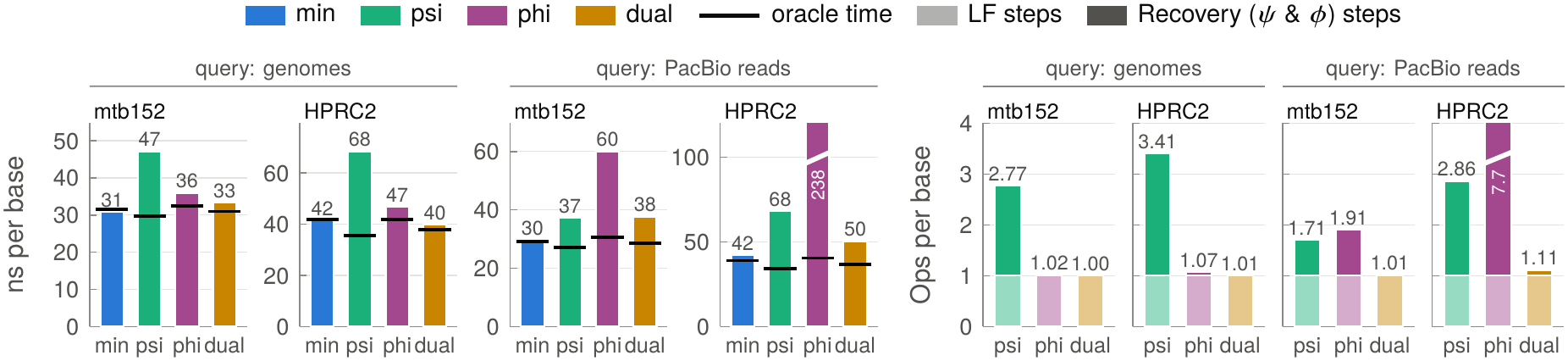
Time (left) and move structure operations (right) per base when performing MS-only queries on mtb152 and HPRC2 pangenomes, querying either 100 kbase pieces of the held-out genomes (“genomes”) or simulated PacBio HiFi reads drawn from them (“PacBio reads”). Queries required 0.00065 recoveries per base for mtb152 genomes, 0.021 for mtb152 PacBio reads, 0.00055 for HPRC2 genomes and 0.036 for HPRC2 PacBio reads. Black ticks mark the time taken by an oracle that uses simple lookups to resolve recoveries. The oracle therefore performs only LF steps (light shading in right panel) whereas the other approaches add further *ϕ* or *Ψ* steps (dark shading). *Ψ* and *ϕ* steps were counted by an instrumented build of the same ms2 version (tag `paper-v1`); LF steps are drawn as one per base rather than counted. ms2-min is omitted from the operations panel since its recovery does not use move structure operations.

For the query reads, we simulated 60 Mbases of PacBio HiFi reads from the genomes. Specifically, we used pbsim3 to simulate 10 noisy passes (subreads), then called consensus reads with PacBio’s ccs tool. The read length distribution was matched to that of the real PacBio reads described below (median 20 kbases), and consensus reads had a mean predicted accuracy of 99.8%.

The query genomes caused relatively few recovery steps: 0.00065 recoveries per base for the mtb152 genome pieces and 0.00055 for the HPRC2 pieces. For the HiFi reads, we observed 0.021 and 0.036 recoveries per base for mtb152 and HPRC2, respectively. We observed that on the HPRC2 genomes, ms2-min’s and ms2-dual’s times stayed within 6% of the oracle baseline (42 and 40 ns per base, against 42 and 38 for their oracles), and ms2-dual performed only 1.01 operations per base. ms2-psi and ms2-phi added *Ψ* or *ϕ* steps at recoveries, since computing an LCE by stepping costs time proportional to the length or BWT distance it covers, reaching 3.41 and 1.07 operations per base and 68 and 47 ns per base on HPRC2. On mtb152, ms2-min was fastest, followed by ms2-dual, ms2-phi and ms2-psi (31, 33, 36 and 47 ns per base). The PacBio reads required 33 to 65 times as many recoveries per base as the genomes, which widened the gaps to the oracle: ms2-min stayed within 9% of its oracle, but ms2-dual ran 32 to 37% slower than its oracle, and ms2-phi ran 5.9 times slower than its oracle on HPRC2. As expected from its match-length-dependent recovery cost, *Ψ* traversal became cheaper as errors shortened matches (Appendix Figure A2A).

### Comparison with MONI/PHONI on chr22

We compared ms2 to existing programs for computing matching statistics: MONI, PHONI, Augmented PHONI (AugPHONI), and ropebwt3. To keep the experiment manageable for all tools, we indexed only chromosome 22: 100 chromosome 22 haplotypes from HPRC2 (Fig. 4; IDs in Appendix Table A2) and, separately, 10 haplotypes (Appendix Figure A1). For the query reads, we simulated 60 Mbases each of three read types from the long arm of chromosome 22 of a haplotype absent from both indexes: APR008 haplotype 1, the HPRC2 query genome of Fig. 3 (34.0 Mbases). We used only the long arm of the chromosome, since short-arm contigs cannot be scaffolded confidently. We simulated short reads with ART (HiSeq 2500 profile, 125 bp). The PacBio HiFi reads were simulated in the same way as for Fig. 3. We simulated Nanopore reads with pbsim3’s standard ONT model with 96% mean accuracy. Long-read lengths were matched to the real distributions of the PacBio and Nanopore sets described below.

**Fig. 4.**
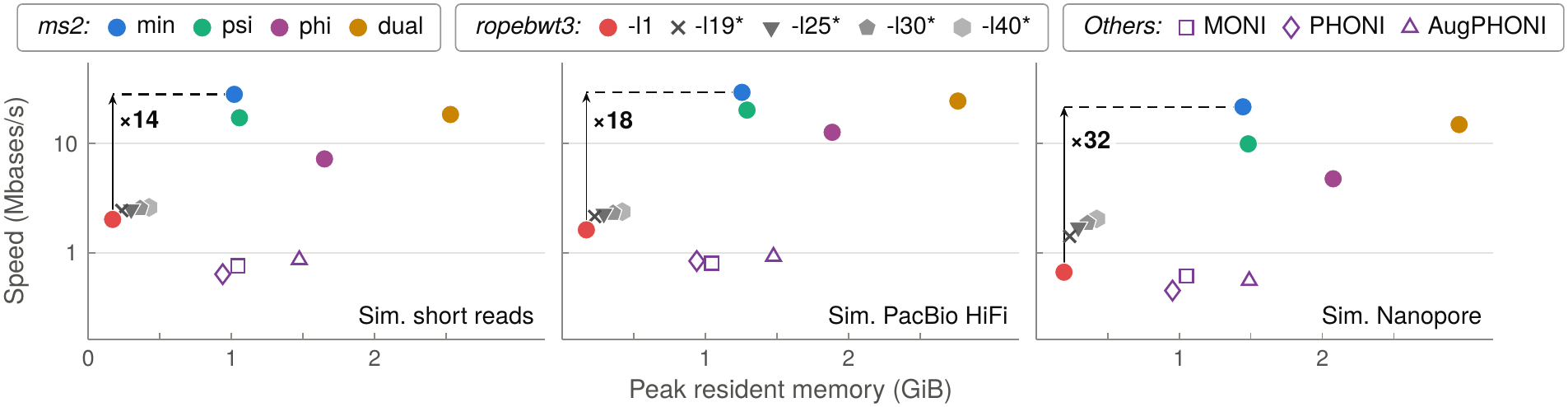
Matching statistics on an index over 100 HPRC2 chromosome 22 haplotypes, queried with simulated reads. ropebwt3 modes marked with an asterisk report only SMEMs of at least the given length, and so have reduced fidelity compared to other modes. Hollow markers indicate where query times were estimated using whole-process time minus time for a one-read run.

With `-l1`, ropebwt3 reports all SMEMs, from which matching statistics can be recovered. With `-l`*L*, how-ever, ropebwt3 reports only SMEMs of length at least *L* (19 by default). We show those faster modes as a baseline, but we mark them with an asterisk to indicate that they are a lower fidelity compared to `-l1` mode, and compared to the other tools. We ran ropebwt3 with `-K 1m`; without that setting, we observed high memory footprints that were due to output buffering. MONI, PHONI and Augmented PHONI do not report timings for index loading and querying phases separately, so we estimated their query time as whole-process time minus that of a run on a single read.

ms2-min computed matching statistics at 22 to 29 Mbases/s, 32 to 35 times faster than the fastest of MONI, PHONI and Augmented PHONI and 35 to 48 times faster than PHONI, with peak memory in the same range (1.0 to 1.4 GiB against 0.9 to 1.5 GiB). ms2-phi was the slowest ms2 mode (4.7 to 12.6 Mbases/s), but it was still 7.7 to 14 times faster than the fastest of MONI, PHONI and Augmented PHONI. ropebwt3 used the least memory by far (0.2 GiB) but computed matching statistics (`-l1`) 14 to 32 times slower than ms2-min. When querying 10 haplotypes instead of 100, the speeds of ms2-min, ms2-psi and ms2-dual changed by 6% or less, while memory shrank with the number of BWT runs (Appendix Figure A1). ms2-phi was the exception: it ran 1.6 to 2.3 times slower on 100 haplotypes than on 10 (where it reached 11 to 20 Mbases/s); a recovery that ms2-phi cannot resolve immediately must use *ϕ* to advance across the run offset-by-offset, and runs tend to lengthen as haplotypes are added (Appendix Figure A2).

### Reads

We queried the HPRC2 index with four sets of real sequencing reads: three from human genomes and one from a skin metagenome. For each, we kept only reads consisting entirely of A, C, G and T (since SMEM-finding tools handle N characters differently), randomly subsampled them, and took whole reads in sampled order up to a total of 60 Mbases; each set holds 59.9 to 60.0 Mbases. **Short reads.** 480,000 Illumina HiSeq 2500 WGS reads (60.0 Mbases) from 2 × 125 bp pairs from the CHM1 hydatidiform mole cell line (SRR3099549). The ropebwt3 study [38] also used this run. CHM1 is not among the HPRC2 haplotypes, so the reads come from a genome absent from the index. Pairs were sampled together, but each end was provided to matching-statistics tools as a separate query. **PacBio.** 2,869 PacBio Sequel II HiFi reads (60.0 Mbases; median length 20.1 kbases) from HPRC sample HG00099 (SRR26545347). Both HG00099 haplotypes are in HPRC2, so this set represents querying reads from a genome in the index. The reads’ base qualities predict a mean error rate of 0.18%. **Nanopore.** 2,706 R10.4.1 reads (59.9 Mbases) from HG002, from the quality-passing reads Oxford Nanopore released in December 2023 (gm24385_2023.12), sampled with `samtools view` at a rate of 0.8% (seed 20261003). Read lengths range from 404 bp to 417 kbases (median 3.4 kbases, N50 91.7 kbases). HG002 is in HPRC2. **Skin metagenome.** 438,969 Illumina NovaSeq 6000 reads (60.0 Mbases; pairs of up to 2 × 151 bp) from an atopic dermatitis skin swab, run ERR14066769 of the hostdepletion study by Guccione et al. [32]. This set represents the screening use case, where most queries do not come from the indexed genomes.

### Matching statistics on real reads

We again compared each ms2 mode with its oracle baseline to assess recovery cost on real reads, using the four real read sets above (Fig. 5). In all four modes, the oracle ran at about 17 to 31 Mbases/s. On human reads, ms2-min ran 1.04 to 1.3 times slower than its oracle, and ms2-psi and ms2-dual ran 1.2 to 2.7 times slower than theirs. The skin metagenome reads necessitated more recoveries and had a wider oracle gap: ms2-min ran at 10.2 Mbases/s versus 26.1 for its oracle. ms2-psi and ms2-dual ran at 2.8 and 5.2 Mbases/s, respectively, versus about 18 for their oracles. ms2-phi was slowest for every read set (0.45 to 7.1 Mbases/s, 16–41% of ms2-psi’s speed) and had the widest gap to its oracle. ms2-min remained 23 to 130 times faster than ropebwt3 computing every matching statistic (`-l1`), which reached 0.08 to 1.12 Mbases/s, and even ms2-phi was 2.6 to 6.3 times faster.

**Fig. 5.**
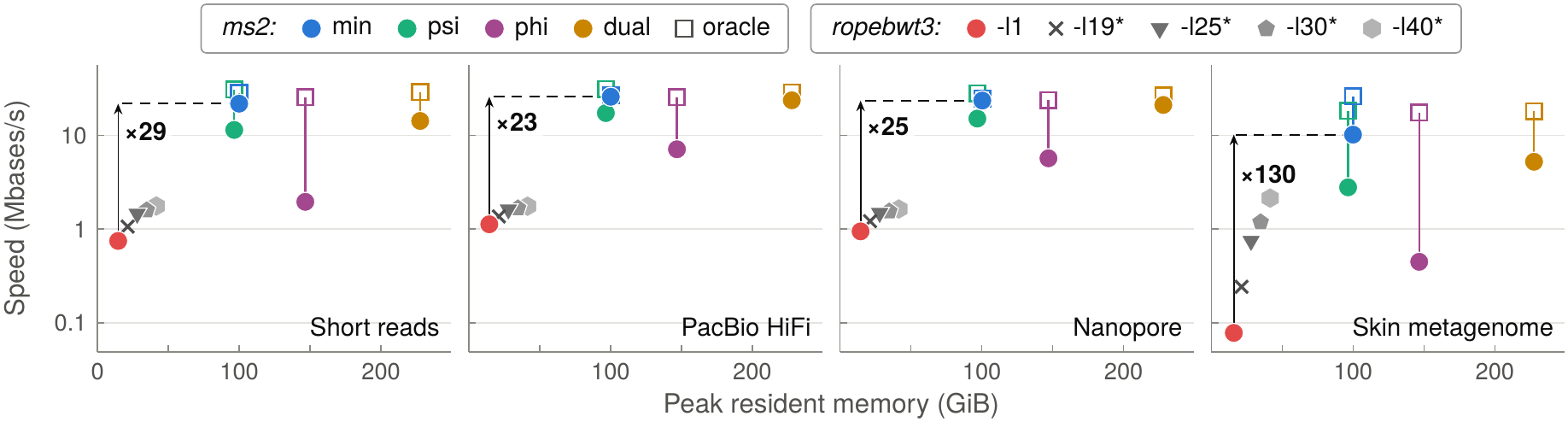
Matching statistics on the HPRC2 index, with oracle replay. For each ms2 approach, a vertical line rises from the ordinary query (circle) to oracle replay (hollow square). Replay points are drawn at the ordinary run’s memory. ropebwt3 modes as in Fig. 4.

To contextualize the speed of these methods, consider that an Oxford Nanopore PromethION R10.4.1 flow cell consists of 2,675 channels [46], each able to sequence 420 bases per second [47]. A flow cell can therefore produce a peak of 1.12 Mbases/s. A single ms2-min thread was able to compute matching statistics and SMEMs at 23.5 Mbases/s; therefore, one ms2-min thread can keep pace with about 23.5*/*1.12 ≈ 21 PromethION flow cells at full speed.

### Locating SMEM occurrences

Finally, we compared SMEM reporting in a common output format, including each SMEM’s query coordinates, full occurrence count, and up to *N* = 100 text positions (Fig. 6). Since these experiments are slower, we used prefixes of the read sets above: the first 6 Mbases of the short reads (48,000 reads), 20 Mbases of the PacBio reads (953 reads), 10 Mbases of the Nanopore reads (494 reads) and 0.5 Mbases of the skin reads (3,698 reads). ms2-psi, ms2-phi and ms2-dual all load the same 337 GiB index, comprising the shared LF*/Ψ* structure together with *ϕ* and *ϕ^−^*^1^ structures and document boundaries, and locate occurrences by *ϕ* and *ϕ^−^*^1^ steps from a sampled offset. Counting and locating procedures are detailed in Appendix Note A5. ropebwt3 (`-l1-p100`) locates occurrences with a sampled suffix array, whose sampling interval (`-s`) is a specified constant. ms2-psi and ms2-dual reported 5.4 to 8.7 million locations/s on every read set, 25 to 34 times faster than ropebwt3 with `-s8` (97 GiB) and 8.8 to 11 times faster than with `-s6`, the densest sampling we tested, which at 345 GiB used more memory than ms2. ms2-phi reported 4.0 to 8.5 million locations/s; it was the slowest ms2 mode on short reads (0.60 times the speed of ms2-dual) and reached 0.79 to 0.98 times ms2-dual’s speed on the other read sets. The sparser `-s11` sampling fit in 25 GiB but reported only 25 to 30 thousand locations/s.

**Fig. 6.**
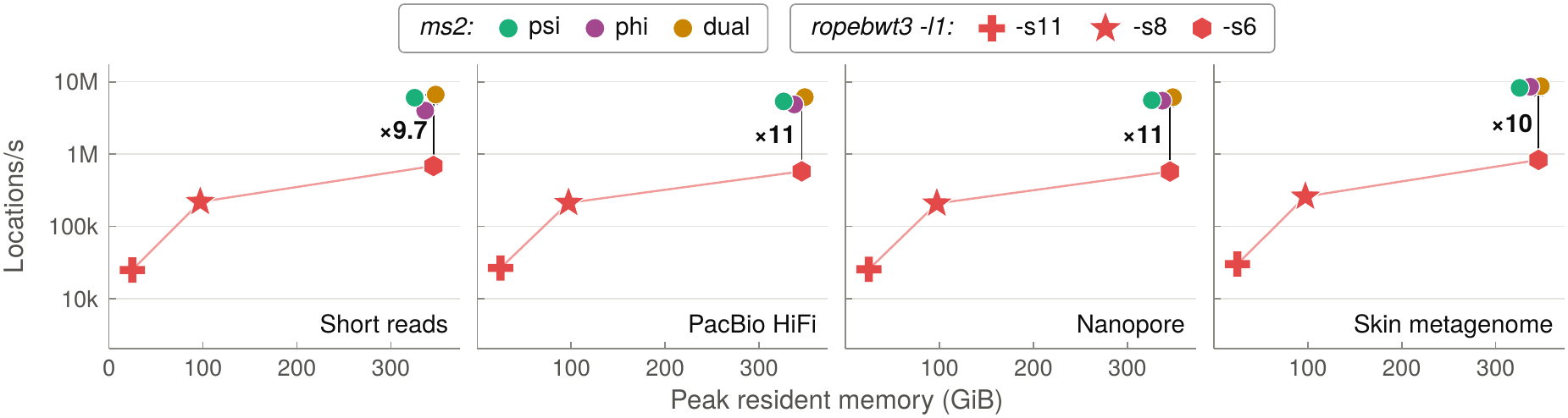
Locating up to *N* = 100 occurrences of every SMEM on the HPRC2 index, against ropebwt3 with suffix-array samples at three sampling intervals. The three ms2 modes load the same index, so their points are offset slightly to the left and right to keep each visible.

### Thread scaling

To measure how the tools scale across threads, we computed matching statistics for 1 Gbase supersets of the short and Nanopore read sets with 1 to 32 threads, one per physical core of a single socket, and with 64 threads using hyperthreading on those 32 cores (Appendix Figure A3). All tools scaled well up to the number of physical cores: with 32 threads, the ms2 modes ran 21.5 to 27.6 times faster than with one thread and ropebwt3 `-l1` ran 25 to 30 times faster. ms2-min reached 469 to 498 Mbases/s. The slowest tools, ms2-phi and ropebwt3, benefited further from hyperthreading, running 1.16 to 1.28 times faster with 64 threads than with 32, while the faster ms2-min and ms2-dual modes did not.

## 5 Discussion

Compressed indexing is crucial for querying pangenomes. To date, no compressed index has been capable of computing matching statistics and SMEMs quickly and in a single pass. We narrowed this gap by lever-aging novel compositions of move structures to make new efficient ways to handle the crucial “recovery” step. In a human pangenome setting, our fastest strategy (ms2-min) approaches the efficiency of an oracle that skips recovery work, indicating that little of the original gap remains. While the methods proposed here are implemented in ms2, they can be adopted by other tools using move structures or *r*-indexes.

When used to compute matching-statistics lengths only, without reporting any SMEM offsets, ms2-min is the fastest of our methods. When additionally reporting SMEM offsets, the best method can vary. In scenarios with shorter match lengths, ms2-psi is favored. When the pangenome consists of fewer sequences and move-structure rows span fewer suffixes each, ms2-phi can be faster. The hybrid ms2-dual mode uses more memory than either ms2-psi or ms2-phi for match length computation, but combines the strengths of the *Ψ* and *ϕ*-based methods, achieving predictably excellent speed across all tested scenarios.

We established query and space bounds for the invertible move structure proposed by Brown et al. [17], which supports both directions of a permutation in a shared table without separate *Ψ* links (Appendix Note A3). Our framework also supports linear-time queries through caching under a BWT-based similarity condition, and occurrence counting by maintaining full SA intervals (Appendix Notes A1 and A2). The latter offers an alternative to forward-backward search for counting and could support tag array queries [28], such as reporting distinct alignment columns associated with a match [18, 45]. We have yet to fully explore the practical benefits of these variants.

In the future, it will be important to reduce the size of the navigation structures, opening new tradeoffs between speed and memory footprint. Combining Boyer-Moore-Li skipping with move structures could accelerate searches restricted to long SMEMs. Suffixient sets [23] and suffix tree path decompositions [9] also merit exploration for matching statistics computation.

These methods provide fast exact lengths and locations for downstream alignment, host screening, and classification. Their throughput brings exact matching on large pangenomes within reach of real-time sequencing workflows.

## Acknowledgments

N.K.B. was supported by an NSERC PGS-D and NIH grant R01HG011392. B.L. was supported by NIH grant R01HG011392. A.S., P.S., and S.Z. were supported by NIH grant R01HG010086. The ms2 code was implemented by N.K.B. and B.L. with the help of Claude Code and OpenAI Codex, though all of that work is based on a reference implementation made by hand by N.K.B., A.S., and P.S. The experiments were driven by software made by N.K.B. and B.L. with the help of Claude Code.

## Appendix

**Figure A1.**
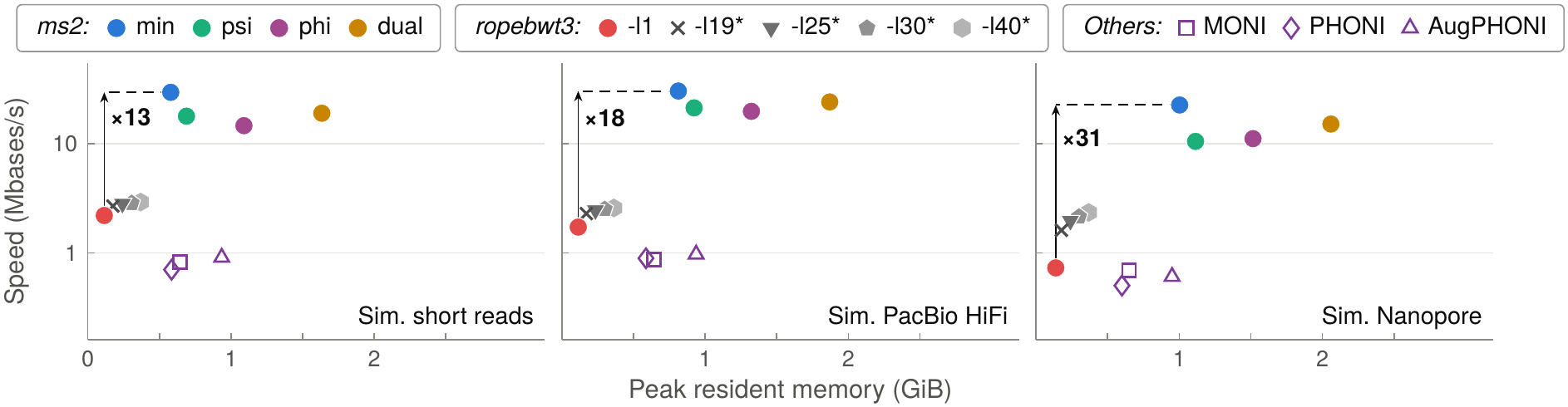
Identical experiment to Fig. 4, but using a chromosome 22 index of 10 HPRC2 haplotypes from 5 donors.

**Figure A2.**
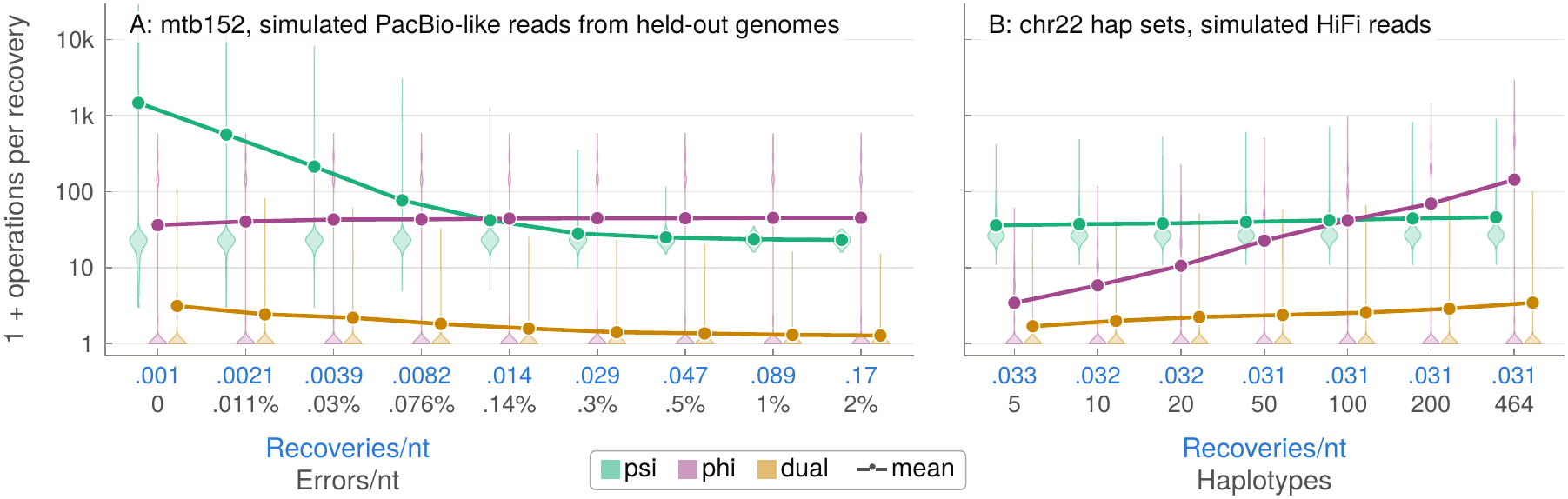
Move structure operations (*Ψ* and *ϕ* steps) per recovery for ms2-psi, ms2-phi and ms2-dual, counted at every recovery by an instrumented build of ms2. Violin plots show each batch’s distribution of one plus the number of operations. Adding 1 was necessary because some recoveries can be resolved without any move structure “operation,” simply by consulting saved LCPs for the current run and surrounding runs (summary skipping). Lines connect the mean values from each batch; the means quoted below count operations without the added one. (A) mtb152 queried with 20 Mbases of PacBio-like reads simulated from the 8 held-out genomes, with independent errors at 0 to 2% per base (bottom tick line). As the recovery rate rose (blue, top tick line) and the lengths of the matches shortened, ms2-psi’s mean fell from 1,471 to 41 at 0.14% errors, where it met ms2-phi’s, and then leveled off near 22, about half of ms2-phi’s, which stayed between 35 and 44. (B) The simulated PacBio HiFi reads of Fig. 4 queried against indexes of the first 5, 10, 20, 50, 100 and 200 HPRC2 chromosome 22 haplotypes and of all 464 (Table A2). As the pangenome grew and the BWT runs lengthened, ms2-phi’s mean rose from 2.4 to 143, meeting ms2-psi’s at 100 haplotypes, while ms2-psi’s stayed between 35 and 45. ms2-dual’s mean stayed between 0.7 and 2.5, well below the mean for both *Ψ* and *ϕ*. This is because dual can stop as soon as either its concurrent *Ψ* or *ϕ* steps resolve the recovery. The cheaper of the two is often far cheaper than the other, as the violins show. Dual also uses lower bounds from *Ψ* comparisons and upper bounds from LCP enumeration to discard a candidate before its exact retained length is known. The batches at 1% and 2% errors and the indexes of 200 and 464 haplotypes were counted on a separate cluster with an instrumented build of the same ms2 version (tag `paper-v1`), outside the benchmark protocol of the other experiments; the other batches come from the benchmark runs.

**Figure A3.**
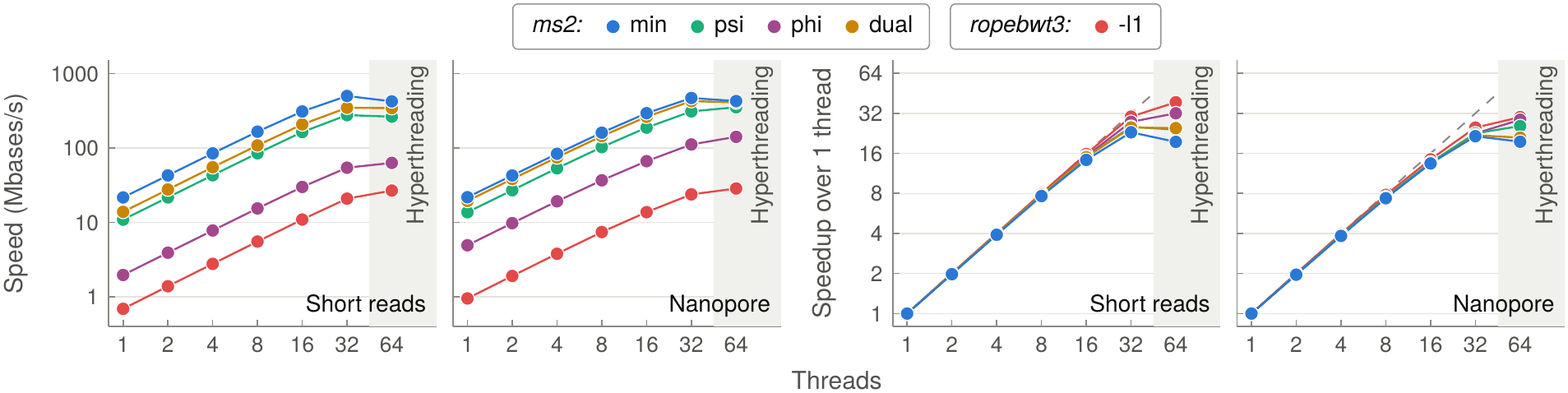
Thread scaling of matching statistics queries on the HPRC2 index. Left: query speed against the number of threads for the four ms2 modes and ropebwt3 `-l1`. Right: speedup relative to one thread; the dashed line marks linear scaling up to 32 threads. Query sets: short reads, 7,982,098 reads, 997.8 Mbases; Nanopore reads, 43,346 reads, 1,000.0 Mbases. The first 60 Mbases of each set are exactly the query set of the same name in Fig. 5. With up to 32 threads, each thread was bound to its own physical core of the first socket. At 64 threads (gray band), two threads shared each of those 32 cores through hyperthreading, also called simultaneous multithreading (SMT). Memory was bound to the same socket in every run. ms2 split the reads into 2 MiB blocks, and ropebwt3 ran with `-t` set to the thread count. Other settings as in Note A6. Note that query interleaving was left at its default of 32 concurrent queries at every thread count, so the query speeds (left) reflect both thread scaling and interleaving.

**Figure A4.**
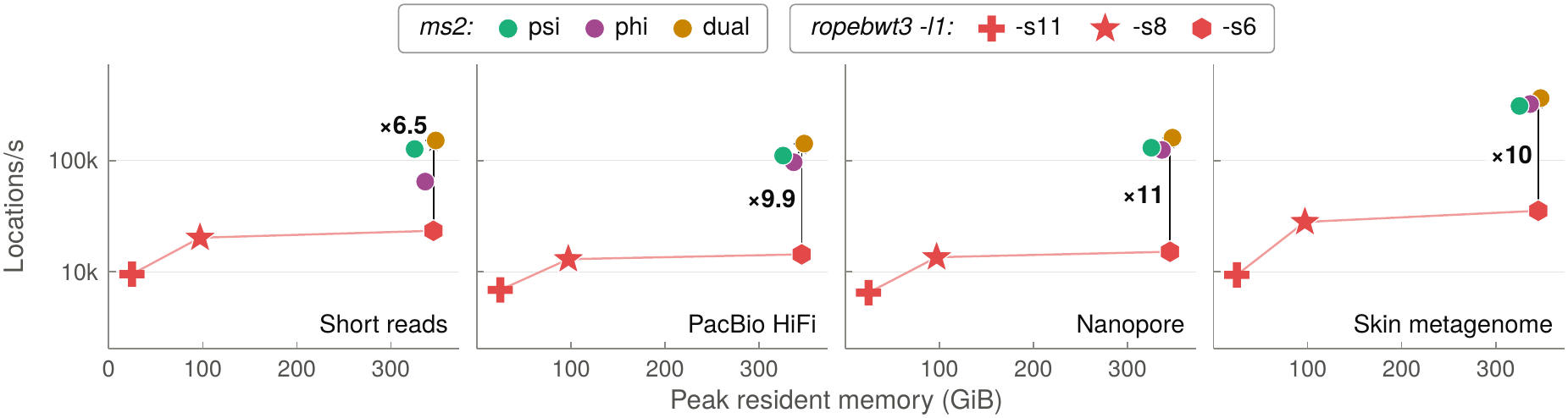
Identical experiment to Fig. 6, but reporting at most *N* = 1 occurrence per SMEM.

**Figure A5.**
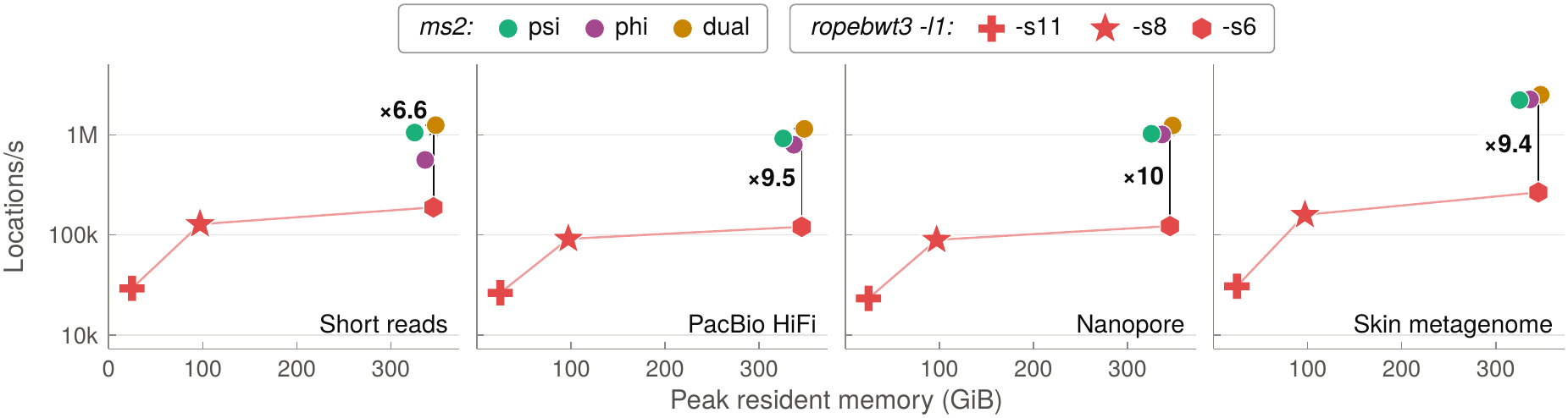
Identical experiment to Fig. 6, but reporting at most *N* = 10 occurrences per SMEM.

**Table A1.**
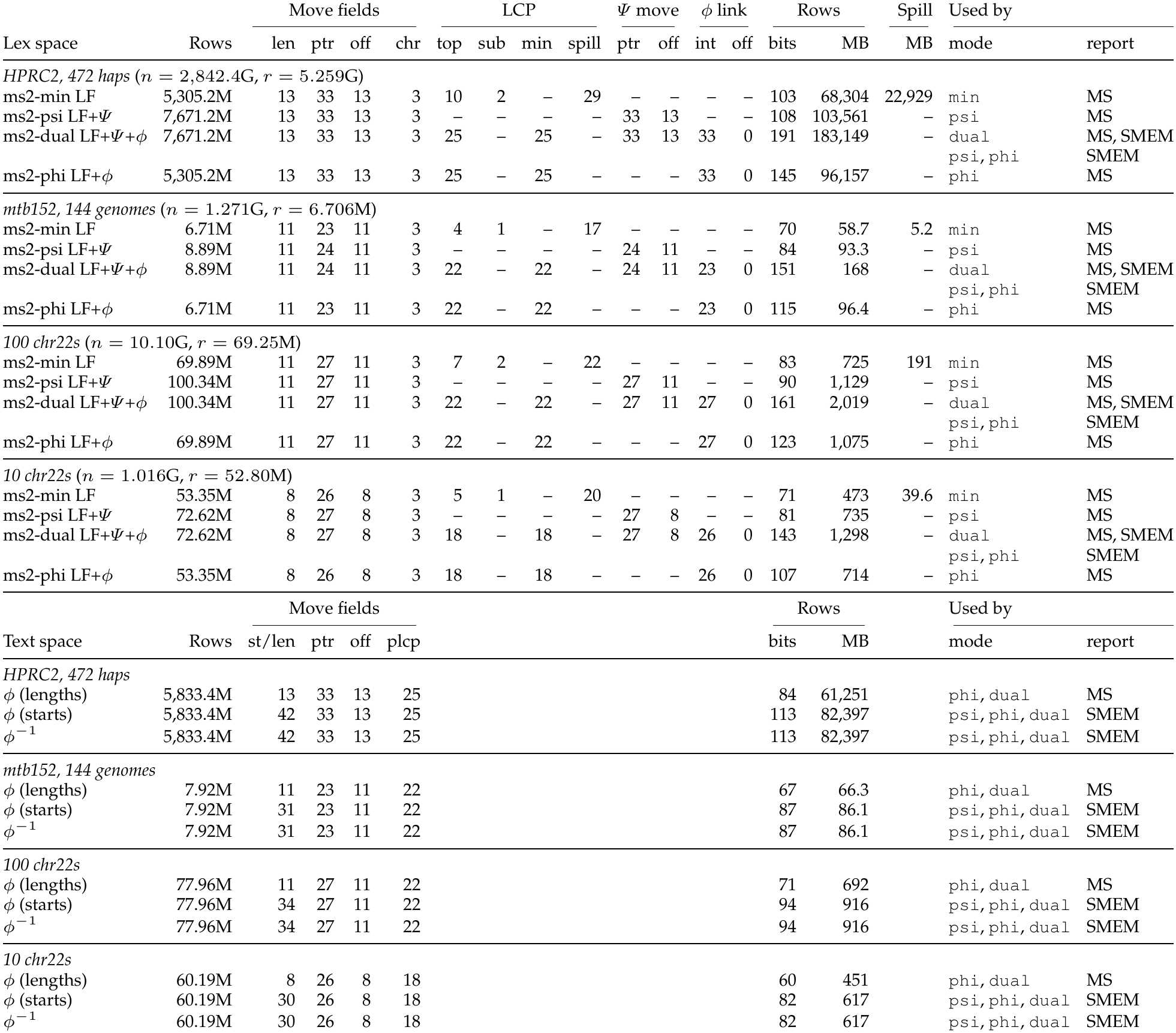
Field widths, in bits, of the move structures for ms2 modes used in our experiments. Rows are in millions (M), and *n* and *r* are the text length and the number of BWT runs. “Lex space” structures are listed in the top section; these are lexicographically ordered structures like the BWT itself, LF and *Ψ*. “Text-space” structures are below; these are structures ordered by text offsets, like *ϕ* and *ϕ^−^*^1^. Basic move structure fields are interval length (len), destination interval (ptr), offset within the destination interval (off) and BWT character (chr). LCP fields can hold each LF interval’s boundary summary (top). For ms2-min, sub stores the difference between the boundary LCP summary and the interval minimum. The spill field points to the interval’s spillover record, holding LCP-minima samples and, in cases where they do not fit in the main table, the full top and minimum LCP values. ms2-min sizes top and sub to the bit width of their 98th percentiles (at least one bit), relying on the spillover for any larger values. ms2-phi and ms2-dual store each interval’s interior minimum (min), with top and min sized to the largest stored value (no spillover). The *Ψ* move fields give the destination interval (ptr) and offset (off) for inverse navigation within the shared LF*/Ψ* structure, mirroring LF move values. *ϕ* links map the interval head to the *ϕ* interval containing its SA value. Text-space rows store either the interval’s start position or its length (st/len) and a sample of the permuted LCP array (plcp). “Rows MB” tallies the space used by the packed rows (rows *×* bits*/*8*/*10^6^), and “Spill MB” gives total size of ms2-min’s spillover arrays. The “Used by” column lists the query and reporting modes that read the structure described in that row. “MS” refers to modes reporting matching statistics only (`--report ms`, the default), and “SMEM” refers to modes that list SMEM occurrences (`--report smem-all`), as in our locating experiments.

| Lex space | Rows | Move fields |  |  |  | LCP |  |  |  | Ψ move |  | ϕ link |  | Rows |  | Spill | Used by |  |
|---|---|---|---|---|---|---|---|---|---|---|---|---|---|---|---|---|---|---|
|  |  | len | ptr | off | chr | top | sub | min | spill | ptr | off | int | off | bits | MB | MB | mode | report |
| *HPRC2, 472 haps* (*n* = 2,842.4G, *r* = 5.259G) |  |  |  |  |  |  |  |  |  |  |  |  |  |  |  |  |  |  |
| ms2-min LF | 5,305.2M | 13 | 33 | 13 | 3 | 10 | 2 | – | 29 | – | – | – | – | 103 | 68,304 | 22,929 | min | MS |
| ms2-psi LF+Ψ | 7,671.2M | 13 | 33 | 13 | 3 | – | – | – | – | 33 | 13 | – | – | 108 | 103,561 | – | psi | MS |
| ms2-dual LF+Ψ+ϕ | 7,671.2M | 13 | 33 | 13 | 3 | 25 | – | 25 | – | 33 | 13 | 33 | 0 | 191 | 183,149 | – | dual | MS, SMEM |
|  |  |  |  |  |  |  |  |  |  |  |  |  |  |  |  |  | psi,phi | SMEM |
| ms2-phi LF+ϕ | 5,305.2M | 13 | 33 | 13 | 3 | 25 | – | 25 | – | – | – | 33 | 0 | 145 | 96,157 | – | phi | MS |
| *mtb152, 144 genomes* (*n* = 1.271G, *r* = 6.706M) |  |  |  |  |  |  |  |  |  |  |  |  |  |  |  |  |  |  |
| ms2-min LF | 6.71M | 11 | 23 | 11 | 3 | 4 | 1 | – | 17 | – | – | – | – | 70 | 58.7 | 5.2 | min | MS |
| ms2-psi LF+Ψ | 8.89M | 11 | 24 | 11 | 3 | – | – | – | – | 24 | 11 | – | – | 84 | 93.3 | – | psi | MS |
| ms2-dual LF+Ψ+ϕ | 8.89M | 11 | 24 | 11 | 3 | 22 | – | 22 | – | 24 | 11 | 23 | 0 | 151 | 168 | – | dual | MS, SMEM |
|  |  |  |  |  |  |  |  |  |  |  |  |  |  |  |  |  | psi,phi | SMEM |
| ms2-phi LF+ϕ | 6.71M | 11 | 23 | 11 | 3 | 22 | – | 22 | – | – | – | 23 | 0 | 115 | 96.4 | – | phi | MS |
| *100 chr22s* (*n* = 10.10G, *r* = 69.25M) |  |  |  |  |  |  |  |  |  |  |  |  |  |  |  |  |  |  |
| ms2-min LF | 69.89M | 11 | 27 | 11 | 3 | 7 | 2 | – | 22 | – | – | – | – | 83 | 725 | 191 | min | MS |
| ms2-psi LF+Ψ | 100.34M | 11 | 27 | 11 | 3 | – | – | – | – | 27 | 11 | – | – | 90 | 1,129 | – | psi | MS |
| ms2-dual LF+Ψ+ϕ | 100.34M | 11 | 27 | 11 | 3 | 22 | – | 22 | – | 27 | 11 | 27 | 0 | 161 | 2,019 | – | dual | MS, SMEM |
|  |  |  |  |  |  |  |  |  |  |  |  |  |  |  |  |  | psi,phi | SMEM |
| ms2-phi LF+ϕ | 69.89M | 11 | 27 | 11 | 3 | 22 | – | 22 | – | – | – | 27 | 0 | 123 | 1,075 | – | phi | MS |
| *10 chr22s* (*n* = 1.016G, *r* = 52.80M) |  |  |  |  |  |  |  |  |  |  |  |  |  |  |  |  |  |  |
| ms2-min LF | 53.35M | 8 | 26 | 8 | 3 | 5 | 1 | – | 20 | – | – | – | – | 71 | 473 | 39.6 | min | MS |
| ms2-psi LF+Ψ | 72.62M | 8 | 27 | 8 | 3 | – | – | – | – | 27 | 8 | – | – | 81 | 735 | – | psi | MS |
| ms2-dual LF+Ψ+ϕ | 72.62M | 8 | 27 | 8 | 3 | 18 | – | 18 | – | 27 | 8 | 26 | 0 | 143 | 1,298 | – | dual | MS, SMEM |
|  |  |  |  |  |  |  |  |  |  |  |  |  |  |  |  |  | psi,phi | SMEM |
| ms2-phi LF+ϕ | 53.35M | 8 | 26 | 8 | 3 | 18 | – | 18 | – | – | – | 26 | 0 | 107 | 714 | – | phi | MS |
|  |  | Move fields |  |  |  |  |  |  |  |  |  |  |  | Rows |  |  | Used by |  |
| Text space | Rows | st/len | ptr | off | plcp |  |  |  |  |  |  |  |  | bits | MB |  | mode | report |
| *HPRC2, 472 haps* |  |  |  |  |  |  |  |  |  |  |  |  |  |  |  |  |  |  |
| ϕ (lengths) | 5,833.4M | 13 | 33 | 13 | 25 |  |  |  |  |  |  |  |  | 84 | 61,251 |  | phi,dual | MS |
| ϕ (starts) | 5,833.4M | 42 | 33 | 13 | 25 |  |  |  |  |  |  |  |  | 113 | 82,397 |  | psi,phi,dual | SMEM |
| ϕ<sup>−1</sup> | 5,833.4M | 42 | 33 | 13 | 25 |  |  |  |  |  |  |  |  | 113 | 82,397 |  | psi,phi,dual | SMEM |
| *mtb152, 144 genomes* |  |  |  |  |  |  |  |  |  |  |  |  |  |  |  |  |  |  |
| ϕ (lengths) | 7.92M | 11 | 23 | 11 | 22 |  |  |  |  |  |  |  |  | 67 | 66.3 |  | phi,dual | MS |
| ϕ (starts) | 7.92M | 31 | 23 | 11 | 22 |  |  |  |  |  |  |  |  | 87 | 86.1 |  | psi,phi,dual | SMEM |
| ϕ<sup>−1</sup> | 7.92M | 31 | 23 | 11 | 22 |  |  |  |  |  |  |  |  | 87 | 86.1 |  | psi,phi,dual | SMEM |
| *100 chr22s* |  |  |  |  |  |  |  |  |  |  |  |  |  |  |  |  |  |  |
| ϕ (lengths) | 77.96M | 11 | 27 | 11 | 22 |  |  |  |  |  |  |  |  | 71 | 692 |  | phi,dual | MS |
| ϕ (starts) | 77.96M | 34 | 27 | 11 | 22 |  |  |  |  |  |  |  |  | 94 | 916 |  | psi,phi,dual | SMEM |
| ϕ<sup>−1</sup> | 77.96M | 34 | 27 | 11 | 22 |  |  |  |  |  |  |  |  | 94 | 916 |  | psi,phi,dual | SMEM |
| *10 chr22s* |  |  |  |  |  |  |  |  |  |  |  |  |  |  |  |  |  |  |
| ϕ (lengths) | 60.19M | 8 | 26 | 8 | 18 |  |  |  |  |  |  |  |  | 60 | 451 |  | phi,dual | MS |
| ϕ (starts) | 60.19M | 30 | 26 | 8 | 18 |  |  |  |  |  |  |  |  | 82 | 617 |  | psi,phi,dual | SMEM |
| ϕ<sup>−1</sup> | 60.19M | 30 | 26 | 8 | 18 |  |  |  |  |  |  |  |  | 82 | 617 |  | psi,phi,dual | SMEM |

**Table A2.** HPRC2 donors in index order, two chromosome 22 haplotypes each (paternal or first, then maternal or second). An index of *h* haplotypes holds the first *h* haplotypes of this list: the 10-haplotype index contains the first 5 donors (bold), the 100-haplotype index contains the first 50 (above the first rule), the 200-haplotype index of Figure A2 contains the first 100 (above the second rule), and the 464-haplotype index contains all 232.

| **HG002** | **HG06807** | **HG00408** | **HG00423** | **HG00438** | HG00544 | HG00558 | HG00597 | HG005 | HG00609 |
|---|---|---|---|---|---|---|---|---|---|
| HG00621 | HG00639 | HG00642 | HG00658 | HG00673 | HG00706 | HG00733 | HG00735 | HG00738 | HG00741 |
| HG01071 | HG01074 | HG01081 | HG01099 | HG01106 | HG01109 | HG01123 | HG01150 | HG01175 | HG01192 |
| HG01243 | HG01252 | HG01255 | HG01258 | HG01261 | HG01346 | HG01358 | HG01361 | HG01433 | HG01496 |
| HG01884 | HG01891 | HG01928 | HG01934 | HG01940 | HG01943 | HG01952 | HG01960 | HG01969 | HG01975 |
| HG01981 | HG01993 | HG02004 | HG02015 | HG02027 | HG02055 | HG02056 | HG02071 | HG02074 | HG02080 |
| HG02083 | HG02109 | HG02129 | HG02132 | HG02135 | HG02145 | HG02148 | HG02258 | HG02280 | HG02293 |
| HG02300 | HG02451 | HG02486 | HG02514 | HG02523 | HG02559 | HG02572 | HG02602 | HG02615 | HG02622 |
| HG02630 | HG02647 | HG02668 | HG02698 | HG02717 | HG02723 | HG02735 | HG02738 | HG02809 | HG02818 |
| HG02841 | HG02886 | HG02984 | HG03017 | HG03041 | HG03050 | HG03098 | HG03239 | HG03453 | HG03486 |
| HG03540 | HG03579 | HG03654 | HG03669 | HG03688 | HG03704 | HG03710 | HG03804 | HG03816 | HG03831 |
| HG03834 | HG03927 | HG03942 | HG04115 | HG04157 | HG04160 | HG04184 | HG04187 | HG04199 | HG04204 |
| HG04228 | NA18906 | NA19240 | NA20129 | NA21309 | HG01978 | HG02257 | HG03516 | HG00097 | HG00099 |
| HG00126 | HG00128 | HG00133 | HG00140 | HG00146 | HG00232 | HG00235 | HG00253 | HG00272 | HG00280 |
| HG00290 | HG00320 | HG00321 | HG00323 | HG00329 | HG00344 | HG00350 | HG01167 | HG01530 | HG01784 |
| HG01786 | HG02040 | HG02155 | HG02165 | HG02178 | HG02273 | HG02391 | HG02392 | HG02583 | HG02922 |
| HG02965 | HG02976 | HG03130 | HG03139 | HG03195 | HG03209 | HG03225 | HG03270 | HG03369 | HG03470 |
| HG03471 | HG03521 | HG03583 | HG03742 | HG03784 | HG03874 | NA18505 | NA18508 | NA18522 | NA18565 |
| NA18570 | NA18608 | NA18620 | NA18747 | NA18879 | NA18940 | NA18943 | NA18944 | NA18945 | NA18948 |
| NA18952 | NA18959 | NA18960 | NA18967 | NA18970 | NA18971 | NA18974 | NA18976 | NA18982 | NA18983 |
| NA19036 | NA19043 | NA19087 | NA19159 | NA19185 | NA19338 | NA19391 | NA19443 | NA19468 | NA19682 |
| NA19700 | NA19776 | NA19835 | NA19909 | NA20282 | NA20346 | NA20503 | NA20752 | NA20762 | NA20799 |
| NA20805 | NA20806 | NA20809 | NA20827 | NA20850 | NA20870 | NA20905 | NA21093 | NA21102 | NA21106 |
| NA21110 | NA21144 |  |  |  |  |  |  |  |  |

**Table A3.**
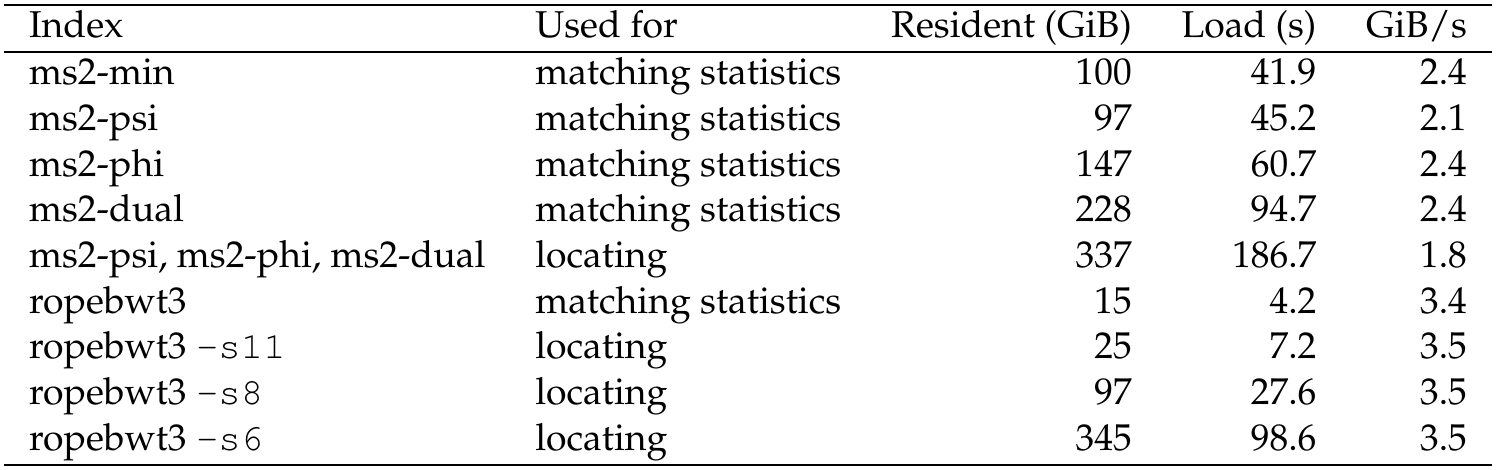
Index load time measurements for HPRC2 indexes, computed when loading from a node-local disk. All times are medians over the same runs used for Figs. 5 and 6. Resident size is the peak memory of a query run.

### Note A1 Matching Similar Patterns

We present an argument for why recovery-style algorithms like the ones described here can compute matching statistics in time linear in the length of the pattern *P* under certain similarity assumptions. The argument uses the average BWT run length, which is one measure of text repetitiveness. Let the terminator already included in *T* be replaced by $_1_, with $_1_ *<* $_2_ *< c* for any *c* ∈ *Σ*. Then let *r_P_* be the number of runs in BWT(*P* $_2_) and *R* the number of runs in the multi-dollar BWT [A2] of *T* [0*..n* − 2]$_1_*P* $_2_ = (*T, P*). We

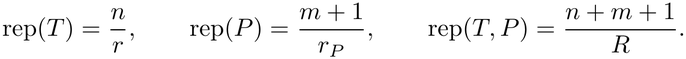

Intuitively, combining two similar strings yields a collection at least as repetitive as either string alone. Therefore, we refer to a pattern *P* as *similar* to *T* if max{rep(*T*), rep(*P*)} ≤ rep(*T, P*).

**Theorem 1.** Given an O(*r*)*-space ms2-phi index, the matching statistics of a pattern P similar to T, whose characters all occur in T, can be computed in O*(*m*)*-time and O*(*r* + *m*) *total space*.

*Proof.* Initializing *p* = 0 and *ℓ* = 0 makes the initial candidate a lexicographic neighbor of $_2_. The candidate remains a neighbor of *P* [*i..m* − 1]$_2_ during match extension and recovery: direct LF extension preserves adjacency, and on a mismatch the nearest preceding and following eligible BWT positions map to neighbors of the extended suffix.

Because $_1_ *<* $_2_ *< c* for any *c* ∈ *Σ*, BWT(*T, P*) is obtained by inserting the pattern rows into gaps of BWT(*T*) (with its terminator relabeled to $_1_). Several recoveries may fall in the same gap or move interval without creating further runs. To avoid charging an interval repeatedly, the first recovery requiring enumeration in interval *j* with range [*s_j_, e_j_*] scans all interior LCP values with *ϕ* and stores the directional minima 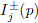 for every interior position in uncompressed arrays of length *e_j_* − *s_j_*. Length capping bounds this size to *O*(*n/r*), resulting in *O*(*n/r*)-time and space construction. Later recoveries at any interior rank *p* look up 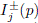 directly via offset *p* − *s_j_* in *O*(1)-time (avoiding the binary search used in ms2-min), while recoveries at interval endpoints already use the existing summaries in constant time.

Let *K* count the cached intervals, i.e., the unique intervals requiring interior enumeration during recovery. Each first enumeration has a candidate strictly inside an interval of BWT character *a* and requests *c* = *a*. By adjacency, the current pattern row inserts *c* between two text *a*’s in that interval, creating at least two new run boundaries. Distinct cached intervals charge distinct gaps, and other insertions cannot remove existing run boundaries, so similarity gives

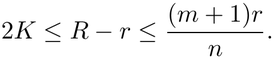

Cache construction therefore takes *O*(*K* · *n/r*) = *O*(*m*)-time and space. All other operations take constant time per character, yielding *O*(*m*) query time and *O*(*r* + *m*) total space.

**Corollary 1.** Given an O(*r* + *S*)*-space ms2-min index, the matching statistics of a pattern P similar to T, whose characters all occur in T, can be computed in O*(*m*)*-time and O*(*r* + *S* + *m*) *total space*.

*Proof.* On the first recovery requiring a directional query in interval *j*, expand its retained directional chains into arrays indexed by interior position. Initialize the running minima to *b_j_* and *b_j_*_+1_, respectively, and fill positions between retained samples with the current directional minimum. The resulting arrays suffice because each recovery range includes the corresponding boundary. Length capping bounds construction by *O*(*n/r*)-time and space, and subsequent queries use constant-time lookup. The preceding charging argument bounds the total construction cost by *O*(*m*).

### Note A2 Maintaining the Suffix Array Interval

We can maintain the full SA interval [*a, b*] of the current match and its length *ℓ*. The interval width gives the occurrence count. We maintain the interval through backward search and, when extension fails, recover it by searching outward from a BWT run boundary inside the recovered match interval.

**Theorem 2.** ms2-min supports matching statistics and SMEM occurrence counts in O(*r*+*S*)*-space, with O*(1)*-time extensions and O*(log *r* + log(*n/r*))*-time recoveries*.

*Proof.* Maintain the move intervals containing *a* and *b*. For *P* [*i* − 1], the per-character links find its first and last occurrences in BWT[*a..b*] in constant time. If they exist, applying LF to these positions gives the extended interval, completing one backward search step. If *P* [*i*−1] does not occur in *T*, set *ℓ* = MS[*i*−1] = 0 and [*a, b*] = [0*, n* − 1], and continue to the next character.

If backward extension fails and *P* [*i* − 1] occurs in *T*, apply match recovery from *a* toward the preceding candidate and from *b* toward the following candidate, selecting *q* with the larger retained length *ℓ_q_*. If *ℓ_q_* = 0, set [*a, b*] = [0*, n* − 1] and proceed directly to backward extension. Otherwise, if *q < a*, then *q* is a run tail, so set *t* = *q* + 1; if *q > b*, then *q* is a run head, so set *t* = *q*. The definition of retained length gives LCP[*t*] ≥ *ℓ_q_*, so this run boundary lies inside the recovered match interval.

The recovered interval is [*u, v* − 1], where *u < t < v* are the nearest positions on either side of *t* with LCP below *ℓ_q_*, taking LCP[*n*] = 0. Starting from *t*, binary search with RMQ over *b_j_* and *µ_j_* finds the nearest summary below *ℓ_q_* in each direction in *O*(log *r*)-time. A boundary summary determines the endpoint directly; an interior summary identifies the interval whose directional chain we search in *O*(log(*n/r*))-time. We use the left-to-right chain when searching rightward and the right-to-left chain when searching leftward. The adjacent boundary value is at least *ℓ_q_*, so every sample below *ℓ_q_* is retained; the existing chains therefore suffice.

We initialize as in match extension, additionally setting [*a, b*] = [0*, n* − 1]. After match recovery, extension by *P* [*i* − 1] gives the new interval and sets *ℓ* = MS[*i* − 1] = *ℓ_q_* + 1. Each interval’s width gives the occurrence count for its matching-statistics entry. Retaining the previous width supplies the count when that entry is identified as an SMEM.

**Corollary 2.** ms2-phi supports matching statistics and SMEM occurrence counts in O(*r*)*-space, with O*(1)*-time extensions and O*(log *r* + *n/r*)*-time recoveries*.

*Proof.* Replace each directional-chain search with *ϕ* enumeration within the identified interval, taking *O*(*n/r*)-time. For a rightward search, enumerate from the interval tail and retain the leftmost LCP value below *ℓ_q_*. The summary search and backward extensions are unchanged.

Our implementation instead uses the counting procedure in Note A5.

### Note A3 Invertible Move Structure

We formalize the *invertible move structure* proposed by Brown et al. [17] and establish its query and space bounds. We use their notation: for a permutation *π* over [0*, n*) with *r* maximal intervals on which *π*(*i* + 1) = *π*(*i*) + 1, let *P* [0*..r^′^*] and *Q*[0*..r^′^*] represent the input and output intervals after simultaneous balancing with parameter *α* ≥ 2, where *r^′^* = *O*(*r*), *P* [*r^′^*] = *Q*[*r^′^*] = *n*, and *π*(*P* [*j*]) = *Q*[*τ* (*j*)]. We store both directions over the common partition *U* = *P* ∪ *Q*. Let *U.* rank(*i*) = max{*j*: *U* [*j*] ≤ *i*} denote the index of the interval containing *i*. Coordinates can therefore serve as input to either direction without links for converting between distinct move structures.

**Lemma 1.** The partition U is balanced with respect to both π and π^−1^.

*Proof.* Every interval of *U* lies within an input interval of *P*, so its image lies within an output interval [*q_s_, q_e_*) of *Q*. By simultaneous balancing, |(*q_s_, q_e_*) ∩ *P* | *<* 2*α*, and no element of *Q* lies strictly inside this interval. Thus, the image contains fewer than 2*α* interior boundaries of *U*. The symmetric argument holds for *π^−^*^1^.

Each interval stores its length or starting position and two bits indicating whether its head belongs to *P*, *Q*, or both. A head *p* ∈ *P* stores the interval identifier *U.* rank(*π*(*p*)); symmetrically, a head *q* ∈ *Q* stores *U.* rank(*π^−^*^1^(*q*)). Both images are heads in *U*, so these pointers require no stored image offsets.

**Theorem 3.** Given simultaneously balanced P, Q, and τ, an invertible move structure can be constructed in O(*r^′^*)*-time and space, supporting move queries in either direction in O*(*α*)*-time without stored image offsets*.

*Proof.* Consider a forward query at position *i* in interval *j* = *U.* rank(*i*). Scan backward to the nearest head *p* ∈ *P*. Every crossed boundary belongs to *Q* and lies strictly inside the input interval containing *i*, so simultaneous balancing bounds this scan by 2*α* − 1 steps. Follow the stored pointer to *h* = *U.* rank(*π*(*p*)) and compute *π*(*i*) = *U* [*h*] + (*i* − *p*). Starting at *h*, scan forward to find *j^′^* = *U.* rank(*π*(*i*)). This scan remains within the output interval corresponding to *p*, whose interior contains no *Q*-boundaries and fewer than 2*α P*-boundaries. It therefore crosses at most 2*α* − 1 boundaries of *U*. Return (*π*(*i*)*, j^′^*). Inverse queries are symmetric. For construction, merge the sorted arrays *P* and *Q*, recording each head’s identifier in *U*. For each *j*, these identifiers give the forward pointer from *P* [*j*] to *Q*[*τ* (*j*)] and the inverse pointer back to *P* [*j*]. Each head is processed a constant number of times, giving *O*(*r^′^*) construction time and space.

Let *s* = |*P* ∩ *Q*| − 1 count shared interval starts and *u* = |*U* | − 1 = 2*r^′^* − *s* be the number of intervals in *U*. We store one identifier per interval, with the membership bits identifying its direction. At shared heads, this field stores the forward identifier; the inverse identifier is stored in an auxiliary array, addressed by constant-time rank in a bitvector marking shared heads [A4]. This representation stores exactly *u* + *s* = 2*r^′^* identifiers. For the following space bound, we use relative coordinates, storing interval lengths and representing each position by its interval and offset.

**Theorem 4.** Let P and Q represent r^′^ intervals after simultaneous balancing and length capping with fixed parameters. An invertible move structure supports move queries in both directions in O(1)*-time using*

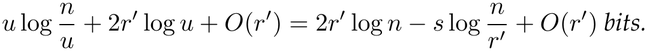

*Proof.* Length capping gives maximum length *O*(*n/u*), so interval lengths occupy *u* log(*n/u*) + *O*(*u*) bits. The 2*r^′^* identifiers occupy 2*r^′^*⌈log *u*⌉ bits, while membership bits and constant-time rank support require *O*(*u*) bits. Accumulating interval lengths replaces absolute differences in the preceding query algorithm, preserving its time bound. Alternatively, a sparse bitvector represents the interval starts within the same space bound and supports constant-time access by interval rank [16]. Finally, *u* = 2*r^′^* − *s* and *r^′^* ≤ *u* ≤ 2*r^′^* give the stated expression.

Two separate length capped structures with *r^′^* intervals each use 2*r^′^* log *r^′^* + 4*r^′^* log(*n/r^′^*) + *O*(*r^′^*) bits [16], excluding additional support for converting between their interval coordinates. The invertible representation saves (2*r^′^* + *s*) log(*n/r^′^*) + *O*(*r^′^*) bits even before accounting for this support. It supports both directions within the leading 2*r^′^* log *n*-bit bound of one classical move structure, while retaining the overall *O*(*r* log *n*)-bit bound.

Orbit implements two offset-free layouts: two identifier fields per interval, or one field with paired overflow records at shared heads. Both retain the backward scan. A third layout stores both directional destinations and one shared offset, avoiding this scan. At a *P* head the forward offset is zero, and at a *Q* head the inverse offset is zero; the membership bits identify which direction uses the stored offset. Our evaluated implementation instead stores separate offsets for both directions. The rank-addressed representation is theoretical and is not used in the evaluated implementation.

### Note A4 Storage Layouts

*Move structure layouts.* Orbit stores move intervals as contiguous bitpacked rows, with fixed column widths and auxiliary data alongside navigation fields [16]. By default, we use length capping factor *c* = 8 and balancing parameter *α* = 16. For a permutation *π*, a standard row stores its interval’s length or absolute start, the interval containing *π*(*s*) for its start *s*, and the corresponding offset. For ms2-psi and ms2-dual, the invertible LF*/Ψ* representation stores both directional interval identifiers and offsets in each shared row. Table A1 gives the field widths of every structure.

*LCP summaries.* For ms2-min, ms2-phi and ms2-dual, we store the LCP value at each original BWT run head. Our splitting procedures introduce new move interval boundaries within these runs. Rather than compute exact LCP values at these boundaries, we store the larger of the minimum LCP values toward the two original run boundaries. Any recovery range crossing a split boundary also extends to an original run boundary, so it already includes a minimum no larger than the stored summary. The added summary therefore leaves the recovery minimum unchanged. During construction, each split interval’s interior summary is the minimum of its retained LCP samples, or a neutral value if none remain. ms2-phi and ms2-dual store these summaries directly in LF rows, in fields wide enough for the largest value (top and min in Table A1). ms2-min also retains directional samples for querying, using the spillover scheme below.

*LCP minima sampling.* Each ms2-min LF interval row is augmented with fields for its boundary summary (`top`), its difference from the interval minimum (`sub`), and a pointer to spillover storage (`spill`). We size `top` and `sub` to the bit widths required by their respective 98th-percentile values. Skinny rows store both values inline; jumbo rows, whose `top` or `sub` exceeds its field, use a reserved marker indicating that the top summary and row minimum reside in spillover storage. The `spill` width accommodates the largest stored pointer. All retained directional samples reside in spillover storage, with both chains merged into one list of interval-relative positions and unnormalized LCP values encoded as variable-length integers (ULEB128). Identical records share storage. Table A1 gives the resulting field widths and spillover sizes.

*Locating structures.* Relative coordinates represent a position by its interval and offset, replacing absolute interval starts with lengths [15]. These suffice for LF*/Ψ* navigation and *ϕ* traversal when computing only match lengths. Locating text offsets instead uses *ϕ* and *ϕ^−^*^1^ move structures with absolute starts; these do not use the invertible representation. Links from LF to *ϕ* map a BWT position *s* to the *ϕ* interval containing SA[*s*] and its offset within that interval. A split LF interval head may require a nonzero offset, but our implementation reads these links only at original BWT run heads, where the offset is zero, so the offset field is omitted (Table A1). Stored document boundaries map each position in the concatenated text to a document identifier and an offset within that document, matching ropebwt3’s output convention.

### Note A5 Query Execution

*Latency hiding.* Each thread processes a batch over a shared index, interleaving up to 32 active queries by default. Orbit splits move queries into starting and finishing steps: we prefetch the destination row and advance other queries before finishing the move. Recovery traversals likewise advance incrementally to overlap memory accesses. When a query finishes, the next pattern in the batch takes its place.

*Recovery scheduling.* We scan upward and downward in the BWT to find recovery candidates, collecting LCP summaries along the way to compute *R*. The scheduler gives each active query a turn, then advances another query to hide memory latency. For ms2-min, a single scan computes the directional minima for both candidates together. Otherwise, on each turn, traversal methods advance both unresolved candidates, preceding first and following second. For each unresolved candidate, a turn performs the following work:

– **ms2-psi:** A *Ψ* move query advances text extraction by one character for comparison with the pattern.
– **ms2-phi:** A *ϕ* move query and LCP sample access update the running LCP minimum.
– **ms2-dual:** A *Ψ* step followed by a *ϕ* step if recovery remains unresolved.

For ms2-dual, enumeration is omitted when its required distance exceeds *ℓ*. In ms2-phi and ms2-dual, following candidate enumeration starts from the next original BWT run head because we do not store text positions at split boundaries. It may therefore cross several move intervals and exceed the theoretical *O*(*n/r*) enumeration bound.

*Recovery stopping conditions.* Let *q_−_* and *q*_+_ denote the preceding and following candidates. For each *q*, we bound its capped retained length by *L_q_* ≤ *ℓ_q_* ≤ *U_q_*. Interval summaries give the upper bound *C_q_* = min{*ℓ, R_q_*}, and *µ_j_* bounds the interior minimum from below. The implementation favors *q_−_* on ties. We summarize recovery stopping conditions:

– **Full-match retention:** If either candidate retains all *ℓ* characters, it can be selected without further evaluating the other candidate.
– **Summary skipping:** In ms2-min, ms2-phi, and ms2-dual, if *C_q_*≤ *µ_j_*, the interior cannot reduce the result, so *ℓ_q_*= *C_q_* without computing *I^±^*(*p*).
– **Extraction stopping:** In ms2-psi and ms2-dual, extraction stops at the first mismatch or after *ℓ* matching characters.
– **Enumeration stopping:** In ms2-phi and ms2-dual, enumeration stops when its running minimum reaches the interior-summary floor *µ_j_*or its range is exhausted.
– **Candidate discarding:** In ms2-phi and ms2-dual, we discard the following candidate when *U_q_*_+_ ≤ *L_q−_*, or the preceding candidate when *U_q−_ < L_q_*_+_. In dual, successful *Ψ* comparisons raise *L_q_*, while LCP enumeration lowers *U_q_*, strengthening candidate discarding. The surviving candidate continues until its capped length is known.

*Counting and locating.* ms2-psi, ms2-phi and ms2-dual support simultaneous computation of matching statistics and locations (i.e. text offsets) of the found SMEMs. When location reporting is enabled, ms2 reports each SMEM’s full occurrence count and the text offsets of the first *N* rows of its BWT interval, where *N* is the smaller of the occurrence count and the user-specified reporting limit. The output fields and format match ropebwt3’s `smem` command.

Starting from the maintained occurrence, we enumerate upward with *ϕ* and downward with *ϕ^−^*^1^ while adjacent LCP values remain at least the match length, counting every occurrence and keeping the positions of the first *N* rows. If enumeration has not finished after the default budget of eight steps plus one quarter of the match length, rounded down, we backward-search the SMEM to obtain its full BWT interval. The interval width gives the count, and a toehold at its first row gives the first position; *N* − 1 further *ϕ^−^*^1^ steps list the rest without LCP tests. For wide intervals, this fallback avoids enumerating all occurrences just to count them, limiting subsequent listing to *N* positions. Enumerations, searches and listings are interleaved across SMEMs to hide memory latency.

### Note A6 Measurement setup

All experiments use `i4i.metal` instances in `us-east-1`. Each instance has two Intel Xeon Platinum 8375C (Ice Lake) processors, with 32 cores per socket, a 2.9 GHz base and 3.5 GHz maximum clock, and 54 MiB of shared L3 cache per socket. 1 TiB of DRAM is split evenly between two NUMA nodes. Note that for “metal” instances, tools run directly on the host without a hypervisor or other contending processes. Each instance has eight local NVMe drives combined with RAID 0, where indexes, read sets and outputs are held. The instances ran Amazon Linux 2023 with Linux 6.1. We set transparent huge pages to `always` for all processes. Compared with an earlier run under the operating system’s default setting, which is `madvise` on Amazon Linux 2023 (that run did not record the setting), this made ms2 1.4 to 2.2 times faster on the HPRC2 index (median 1.8 when computing matching statistics and 1.6 when locating) and ropebwt3 about 1.2 times faster, without changing peak memory.

Except in the thread-scaling experiment (Fig. A3), each run used one thread and was bound by `numactl` to the cores and memory of NUMA node 0. We dropped the operating system’s page cache before each run, so each run has a “cold” start. We ran each tool three times and reported median-of-3 measurements.

We measured index loading time and query time separately. Query time covers reading, parsing, querying and formatting output. For ms2 we used its built-in timers. For ropebwt3 measurements were derived from timestamps in its log. MONI, PHONI and AugPHONI do not report on the phases separately, so we measured query time as the wall time of a full run minus the wall time of a run on a single read. We measured peak memory as maximum resident set size with GNU `time`.

All tools were built from source: ms2 at tag `paper-v1`, ropebwt3 at commit b408339, MONI at release v0.2.2, PHONI at commit 1b81e46 and AugPHONI at commit 7439580. PHONI and AugPHONI included the fixes described in Note A9.

### Note A7 Code availability

The ms2 software used in this paper is archived at https://doi.org/10.5281/zenodo.24044152 (tag `paper-v1`, with its Orbit and TeraTools submodules). The benchmark workflow, run records and figure sources are archived at https://doi.org/10.5281/zenodo.24045318 (ms2-experiments tag `paper-v1.1`).

### Note A8 Data availability

The HPRC2 sequences and Li’s ropebwt3 index of them are available from Zenodo [A3]. We used that record’s sampled suffix array (shift 11) and sequence-length file unchanged, and converted its `human472.fmr` index to the `human472.fmd` index we used with the command

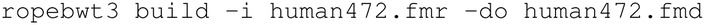

The mtb152 genomes come from Li’s ropebwt3 index collection [39].

We deposited the remaining inputs in three Zenodo records.

– Every read set, with its matching-statistics truth, read offsets and single-read file: https://doi.org/10.5281/zenodo.23964218
– The chromosome 22 and mtb152 RLBWT runs, LCP minima and *ϕ* samples, and the chromosome 22 and mtb152 ropebwt3 indexes: https://doi.org/10.5281/zenodo.23964220
– The MONI, PHONI and Augmented PHONI chromosome 22 indexes, including the indexed texts: https://doi.org/10.5281/zenodo.23964222

Two large HPRC2 inputs are not deposited and must be rebuilt from the `human472.fmd` index. The RLBWT runs, LCP minima and *ϕ* samples (147 GiB) are produced by Orbit’s `fmd2rlbwt` followed by TeraLCP, which took about 13 hours with 48 threads and 171 GiB of memory. The ropebwt3 sampled suffix arrays with shifts 8 and 6 (83 and 331 GiB) are produced by `ropebwt3 ssa-s8` and `-s6`, each taking about 10 hours with 46 to 48 threads. The ms2-experiments repository records the sha256 of every input, so a rebuilt file can be checked before use.

### Note A9 Corrections to PHONI and Augmented PHONI

While validating the matching-statistics outputs of each tool, we found two issues that caused PHONI and Augmented PHONI to report incorrect output. Both tools would sometimes start a query from a BWT row and a reference pointer that referred to different suffixes. Augmented PHONI would additionally read an uninitialized variable in its query loop. The versions we tested include fixes that can be found in pull requests to PHONI (https://github.com/koeppl/phoni/pull/4) and Augmented PHONI (https://github.com/drnatebrown/aug_phoni/pull/2). After these fixes, all tools agree (including with an oracle) on every test query. ropebwt3 was used unmodified. MONI was used unmodified apart from two changes needed to compile it with GCC 13: an added `#include <array>` and a forced include of `cstdint`. PHONI and AugPHONI were also compiled with a forced include of `cstdint` and with `-ftrivial-auto-var-init=zero`, which zero-initializes local variables; without it, PHONI’s index builder can read uninitialized memory and write an invalid index.

### Note A10 Chromosome 22 haplotypes

The chromosome 22 indexes of Fig. 4 and Figure A1 contain chromosome 22 of both haplotypes of the first 50 and the first 5 HPRC2 donors (Table A2), taken in the order of the HPRC release 2 assembly index and excluding CHM13 and GRCh38. The indexes of Figure A2(B) extend the same order to every donor of that index (Table A2); the donor of the simulated reads, APR008, is not among them. Each haplotype’s chromosome 22 sequence was identified by its RagTag [A1] chromosome assignment.

### Note A11 Chromosome 22 with 10 haplotypes

Figure A1 repeats the chromosome 22 comparison of Fig. 4 on an index of 10 HPRC2 haplotypes, queried with the same simulated reads. ms2-min computed matching statistics at 23 to 30 Mbases/s, within 6% of its speed on the 100-haplotype index, while MONI, PHONI and Augmented PHONI ran at 0.5 to 1.0 Mbases/s. Every tool’s peak memory was smaller than on the 100-haplotype index, since the index has fewer BWT runs: 0.6 to 1.0 GiB for ms2-min and 0.6 to 1.0 GiB for MONI, PHONI and Augmented PHONI.

### Note A12 Locating with *N* = 1 and *N* = 10

Figures A4 and A5 repeat the experiment of Fig. 6, reporting at most *N* = 1 and *N* = 10 occurrences per SMEM. With fewer occurrences to report per SMEM, finding the SMEMs was a larger share of the work, and every tool reported fewer locations per second. ms2-psi and ms2-dual remained 5 to 11 times faster than ropebwt3 with `-s6` at both *N* = 1 (111 to 364 thousand locations/s) and *N* = 10 (0.92 to 2.52 million locations/s). As at *N* = 100, ms2-phi was the slowest ms2 mode on short reads (0.43 and 0.45 times the speed of ms2-dual at *N* = 1 and *N* = 10), but it reached 0.68 to 0.90 times ms2-dual’s speed on the other read sets and matched ms2-psi on the Nanopore and skin reads.

### Note A13 Index loading time

For each HPRC2 index, load time grew roughly in proportion to the memory the index occupies once loaded: ms2 loaded at 1.8 to 2.4 GiB/s and ropebwt3 at about 3.5 GiB/s (Table A3). For the 60 Mbases of short reads, loading took 9 to 22 times longer than computing matching statistics with ms2-min, ms2-psi and ms2-dual (and 2 times longer with ms2-phi), so loading dominated the run time of a single query batch of this size.

## References

1. Abeliuk, A., Navarro, G.: Compressed suffix trees for repetitive texts. In: String Processing and Information Retrieval (SPIRE 2012). Lecture Notes in Computer Science, vol. 7608, pp. 30–41. Springer (2012). 10.1007/978-3-642-34109-0_5

2. Abouelhoda, M.I., Kurtz, S., Ohlebusch, E.: Replacing suffix trees with enhanced suffix arrays. Journal of Discrete Algorithms 2(1), 53–86 (2004). 10.1016/S1570-8667(03)00065-0

3. Aganezov, S., Yan, S.M., Soto, D.C., Kirsche, M., Zarate, S., Avdeyev, P., Taylor, D.J., Shafin, K., Shumate, A., Xiao, C., et al.: A complete reference genome improves analysis of human genetic variation. Science 376(6588), eabl3533 (2022). 10.1126/science.abl3533

4. Ahmed, O.Y., Rossi, M., Gagie, T., Boucher, C., Langmead, B.: SPUMONI 2: improved classification using a pangenome index of minimizer digests. Genome Biology 24(1), 122 (2023). 10.1186/s13059-023-02958-1

5. Ahmed, O.Y., Rossi, M., Kovaka, S., Schatz, M.C., Gagie, T., Boucher, C., Langmead, B.: Pangenomic matching statistics for targeted nanopore sequencing. iScience 24(6), 102696 (2021). 10.1016/j.isci.2021.102696

6. Alanko, J.N., Ranallo-Benavidez, T.R., Barthel, F.P., Puglisi, S.J., Marchet, C.: Hierarchical genomic feature annotation with variable-length queries. bioRxiv (2026). 10.64898/2026.03.15.711907

7. Arakawa, Y., Navarro, G., Sadakane, K.: Bi-directional r-indexes. In: 33rd Annual Symposium on Combinatorial Pattern Matching (CPM 2022). Leibniz International Proceedings in Informatics (LIPIcs), vol. 223, pp. 11:1–11:14. Schloss Dagstuhl–Leibniz-Zentrum für Informatik (2022). 10.4230/LIPIcs.CPM.2022.11

8. Bannai, H., Gagie, T., I, T.: Refining the r-index. Theoretical Computer Science 812, 96–108 (2020). 10.1016/j.tcs.2019.08.005

9. Becker, R., Cenzato, D., Gagie, T., Groot Koerkamp, R., Kim, S.H., Manzini, G., Prezza, N.: Compressing suffix trees by path decompositions. In: 53rd International Colloquium on Automata, Languages, and Programming (ICALP 2026). Leibniz International Proceedings in Informatics (LIPIcs), vol. 374, pp. 24:1–24:25. Schloss Dagstuhl–Leibniz-Zentrum für Informatik (2026). 10.4230/LIPIcs.ICALP.2026.24

10. Bender, M.A., Farach-Colton, M.: The LCA problem revisited. In: LATIN 2000: Theoretical Informatics. Lecture Notes in Computer Science, vol. 1776, pp. 88–94. Springer (2000). 10.1007/10719839_9

11. Bertram, N., Fischer, J., Nalbach, L.: Move-r: Optimizing the r-index. In: 22nd International Symposium on Experimental Algorithms (SEA 2024). Leibniz International Proceedings in Informatics (LIPIcs), vol. 301, pp. 1:1–1:19. Schloss Dagstuhl–Leibniz-Zentrum für Informatik (2024). 10.4230/LIPIcs.SEA.2024.1

12. Boucher, C., Gagie, T., I, T., Köppl, D., Langmead, B., Manzini, G., Navarro, G., Pacheco, A., Rossi, M.: PHONI: Streamed matching statistics with multi-genome references. In: 2021 Data Compression Conference (DCC). pp. 193–202. IEEE (2021). 10.1109/DCC50243.2021.00027

13. Boucher, C., Gagie, T., Kuhnle, A., Langmead, B., Manzini, G., Mun, T.: Prefix-free parsing for building big BWTs. Algorithms for Molecular Biology 14(1), 13 (2019). 10.1186/s13015-019-0148-5

14. Brown, N.K., Depuydt, L., Zakeri, M., Alhadi, A., Allam, N., Begleiter, D., Kabilan Karpagavalli, N.B., Khajjayam, S.S., Wahed, H., Gagie, T., Langmead, B.: KeBaB: k-mer based breaking for finding long MEMs. In: String Processing and Information Retrieval (SPIRE 2025). Lecture Notes in Computer Science, vol. 16073, pp. 10–17. Springer (2025). 10.1007/978-3-032-05228-5_2

15. Brown, N.K., Gagie, T., Rossi, M.: RLBWT tricks. In: 20th International Symposium on Experimental Algorithms (SEA 2022). Leibniz International Proceedings in Informatics (LIPIcs), vol. 233, pp. 16:1–16:16. Schloss Dagstuhl– Leibniz-Zentrum für Informatik (2022). 10.4230/LIPIcs.SEA.2022.16

16. Brown, N.K., Langmead, B.: Bounding the average move structure query for faster and smaller RLBWT permutations. In: 24th International Symposium on Experimental Algorithms (SEA 2026). Leibniz International Proceedings in Informatics (LIPIcs), vol. 371, pp. 9:1–9:15. Schloss Dagstuhl–Leibniz-Zentrum für Informatik (2026). 10.4230/LIPIcs.SEA.2026.9

17. Brown, N.K., Sanaullah, A., Zhang, S., Langmead, B.: Optimal-time move structure construction. arXiv preprint arXiv:2603.22147 (2026). 10.48550/arXiv.2603.22147

18. Brown, N.K., Shivakumar, V.S., Langmead, B.: Col-BWT: Pangenomic seed chaining with maximal matches improves read classification. Journal of Computational Biology 33(1), 168–183 (2026). 10.1177/15578666251400784

19. Burrows, M., Wheeler, D.J.: A block-sorting lossless data compression algorithm. Tech. Rep. 124, Digital Equipment Corporation, Systems Research Center (1994)

20. Chang, W.I., Lawler, E.L.: Sublinear approximate string matching and biological applications. Algorithmica 12(4–5), 327–344 (1994). 10.1007/BF01185431

21. Cole, S.T., Brosch, R., Parkhill, J., Garnier, T., Churcher, C., Harris, D., Gordon, S.V., Eiglmeier, K., Gas, S., Barry III, C.E., et al.: Deciphering the biology of *Mycobacterium tuberculosis* from the complete genome sequence. Nature 393(6685), 537–544 (1998). 10.1038/31159

22. Depuydt, L., Ahmed, O.Y., Fostier, J., Langmead, B., Gagie, T.: Run-length compressed metagenomic read classification with SMEM-finding and tagging. iScience 28(12), 114029 (2025). 10.1016/j.isci.2025.114029

23. Depuydt, L., Gagie, T., Langmead, B., Manzini, G., Prezza, N.: Suffixient sets. arXiv preprint arXiv:2312.01359 (2023). 10.48550/arXiv.2312.01359

24. Depuydt, L., Renders, L., Van de Vyver, S., Veys, L., Gagie, T., Fostier, J.: b-move: faster lossless approximate pattern matching in a run-length compressed index. Algorithms for Molecular Biology 20(1), 15 (2025). 10.1186/s13015-025-00281-x

25. Fischer, J., Heun, V.: Space-efficient preprocessing schemes for range minimum queries on static arrays. SIAM Journal on Computing 40(2), 465–492 (2011). 10.1137/090779759

26. Fischer, J., Nalbach, L.: Move-rb: Faster bi-directional r-indexes and approximate pattern matching. arXiv preprint arXiv:2609.30089 (2026). 10.48550/arXiv.2609.30089

27. Gagie, T.: How to find long maximal exact matches and ignore short ones. In: Developments in Language Theory (DLT 2024). Lecture Notes in Computer Science, vol. 14791, pp. 131–140. Springer (2024). 10.1007/978-3-031-66159-4_10

28. Gagie, T.: Tag arrays. arXiv preprint arXiv:2411.15291 (2024). 10.48550/arXiv.2411.15291

29. Gagie, T., Navarro, G., Prezza, N.: Optimal-time text indexing in BWT-runs bounded space. In: Proceedings of the Twenty-Ninth Annual ACM-SIAM Symposium on Discrete Algorithms. pp. 1459–1477. SIAM (2018). 10.1137/1.9781611975031.96

30. Gagie, T., Navarro, G., Prezza, N.: Fully functional suffix trees and optimal text searching in BWT-runs bounded space. Journal of the ACM 67(1), 2:1–2:54 (2020). 10.1145/3375890

31. Goga, A., Depuydt, L., Brown, N.K., Fostier, J., Gagie, T., Navarro, G.: Faster maximal exact matches with lazy LCP evaluation. In: 2024 Data Compression Conference (DCC). pp. 123–132. IEEE (2024). 10.1109/DCC58796.2024.00020

32. Guccione, C., Patel, L., Tomofuji, Y., McDonald, D., Gonzalez, A., Sepich-Poore, G.D., Sonehara, K., Zakeri, M., Chen, Y., Dilmore, A.H., et al.: Incomplete human reference genomes can drive false sex biases and expose patient-identifying information in metagenomic data. Nature Communications 16(1), 825 (2025). 10.1038/s41467-025-56077-5

33. He, Y., Chu, Y., Guo, S., Hu, J., Li, R., Zheng, Y., Ma, X., Du, Z., Zhao, L., Yu, W., et al.: T2T-YAO: a telomere-to-telomere assembled diploid reference genome for Han Chinese. Genomics, Proteomics & Bioinformatics 21(6), 1085–1100 (2023). 10.1016/j.gpb.2023.08.001

34. Hwang, S., Brown, N.K., Ahmed, O.Y., Jenike, K.M., Kovaka, S., Schatz, M.C., Langmead, B.: MEM-based pangenome indexing for *k*-mer queries. Algorithms for Molecular Biology 20, 3 (2025). 10.1186/s13015-025-00272-y

35. Kimura, K., I, T.: R-Enum revisited: Speedup and extension for context-sensitive repeats and net frequencies. In: 37th Annual Symposium on Combinatorial Pattern Matching (CPM 2026). Leibniz International Proceedings in Informatics (LIPIcs), vol. 369, pp. 10:1–10:14. Schloss Dagstuhl–Leibniz-Zentrum für Informatik (2026). 10.4230/LIPIcs.CPM.2026.10

36. Kulmanov, M., Tawfiq, R., Liu, Y., Al Ali, H., Abdelhakim, M., Alarawi, M., Aldakhil, H., Alhattab, D., Alsolme, E.A., Althagafi, A., et al.: A reference quality, fully annotated diploid genome from a Saudi individual. Scientific Data 11(1), 1278 (2024). 10.1038/s41597-024-04121-2

37. Li, H.: Exploring single-sample SNP and INDEL calling with whole-genome de novo assembly. Bioinformatics 28(14), 1838–1844 (2012). 10.1093/bioinformatics/bts280

38. Li, H.: BWT construction and search at the terabase scale. Bioinformatics 40(12), btae717 (2024). 10.1093/bioinformatics/btae717

39. Li, H.: ropebwt indices. Zenodo (2024). 10.5281/zenodo.13147120

40. Lucas, J.K., Hebbar, P., Liao, W.W., Macias-Velasco, J.F., Novak, A.M., Asri, M., Balacco, J.R., Blair, A.P., Ebler, J., Gardner, J.M., et al.: HPRC2: A human pangenome reference with near-complete coverage of common genetic variation. bioRxiv (2026). 10.64898/2026.07.21.739710

41. Marin, M.G., Quinones-Olvera, N., Jin, H., Harris, M.A., Jeffrey, B.M., Rosenthal, A., Murphy, K.C., Sassetti, C.M., Li, H., Farhat, M.R.: Gene conversion is a key driver of diversity hotspots in *Mycobacterium tuberculosis* antigens and virulence-associated loci. Proceedings of the National Academy of Sciences 123(43), e2607261123 (2026). 10.1073/pnas.2607261123

42. Martínez-Guardiola, C., Brown, N.K., Silva-Coira, F., Köppl, D., Gagie, T., Ladra, S.: Augmented thresholds for MONI. In: 2023 Data Compression Conference (DCC). pp. 268–277. IEEE (2023). 10.1109/DCC55655.2023.00035

43. Nishimoto, T., Tabei, Y.: Optimal-time queries on BWT-runs compressed indexes. In: 48th International Colloquium on Automata, Languages, and Programming (ICALP 2021). Leibniz International Proceedings in Informatics (LIPIcs), vol. 198, pp. 101:1–101:15. Schloss Dagstuhl–Leibniz-Zentrum für Informatik (2021). 10.4230/LIPIcs.ICALP.2021.101

44. Ohlebusch, E., Gog, S., Kügel, A.: Computing matching statistics and maximal exact matches on compressed full-text indexes. In: String Processing and Information Retrieval (SPIRE 2010). Lecture Notes in Computer Science, vol. 6393, pp. 347–358. Springer (2010). 10.1007/978-3-642-16321-0_36

45. Olbrich, J., Ohlebusch, E.: The TAG array of a multiple sequence alignment. In: 37th Annual Symposium on Combinatorial Pattern Matching (CPM 2026). Leibniz International Proceedings in Informatics (LIPIcs), vol. 369, pp. 29:1– 29:14. Schloss Dagstuhl–Leibniz-Zentrum für Informatik (2026). 10.4230/LIPIcs.CPM.2026.29

46. Oxford Nanopore Technologies: Flow cells. https://store.nanoporetech.com/flow-cells.html (2026), accessed 2 November 2026

47. Oxford Nanopore Technologies: PromethION 2 integrated technical specification. https://nanoporetech.com/document/requirements/promethion-2i-spec (2026), accessed 2 November 2026

48. Renders, L., Depuydt, L., Gagie, T., Fostier, J.: Columba: fast approximate pattern matching with optimized search schemes. Bioinformatics 41(12), btaf652 (2025). 10.1093/bioinformatics/btaf652

49. Rossi, M., Oliva, M., Langmead, B., Gagie, T., Boucher, C.: MONI: a pangenomic index for finding maximal exact matches. Journal of Computational Biology 29(2), 169–187 (2022). 10.1089/cmb.2021.0290

50. Sanaullah, A., Brown, N.K., Shakya, P., Deegutla, A., Naseri, A., Langmead, B., Zhi, D., Zhang, S.: RLBWT-based LCP computation in compressed space for terabase-scale pangenome analysis. Bioinformatics 42(Supplement_1), btag268 (2026). 10.1093/bioinformatics/btag268

51. Sanaullah, A., Zhi, D., Zhang, S.: An efficient data structure and algorithm for long-match query in run-length compressed BWT. In: 25th International Conference on Algorithms for Bioinformatics (WABI 2025). Leibniz International Proceedings in Informatics (LIPIcs), vol. 344, pp. 17:1–17:25. Schloss Dagstuhl–Leibniz-Zentrum für Informatik (2025). 10.4230/LIPIcs.WABI.2025.17

52. Schneider, V.A., Graves-Lindsay, T., Howe, K., Bouk, N., Chen, H.C., Kitts, P.A., Murphy, T.D., Pruitt, K.D., Thibaud-Nissen, F., Albracht, D., et al.: Evaluation of GRCh38 and de novo haploid genome assemblies demonstrates the enduring quality of the reference assembly. Genome Research 27(5), 849–864 (2017). 10.1101/gr.213611.116

53. Tatarnikov, I., Shahrabi Farahani, A., Kashgouli, S., Gagie, T.: MONI can find k-MEMs. In: 34th Annual Symposium on Combinatorial Pattern Matching (CPM 2023). Leibniz International Proceedings in Informatics (LIPIcs), vol. 259, pp. 26:1–26:14. Schloss Dagstuhl–Leibniz-Zentrum für Informatik (2023). 10.4230/LIPIcs.CPM.2023.26

54. Varki, R., Rossi, M., Ferro, E., Oliva, M., Garrison, E., Langmead, B., Boucher, C.: Accurate short-read alignment through r-index-based pangenome indexing. Genome Research 35(7), 1609–1620 (2025). 10.1101/gr.279858.124

55. Yang, C., Zhou, Y., Song, Y., Wu, D., Zeng, Y., Nie, L., Liu, P., Zhang, S., Chen, G., Xu, J., et al.: The complete and fully-phased diploid genome of a male Han Chinese. Cell Research 33(10), 745–761 (2023). 10.1038/s41422-023-00849-5

56. Zakeri, M., Brown, N.K., Ahmed, O.Y., Gagie, T., Langmead, B.: Movi: a fast and cache-efficient full-text pangenome index. iScience 27(12), 111464 (2024). 10.1016/j.isci.2024.111464

57. Zakeri, M., Brown, N.K., Gagie, T., Langmead, B.: Movi 2: fast and space-efficient queries on pangenomes. Bioinformatics 42(7), btag362 (2026). 10.1093/bioinformatics/btag362

## Appendix References

A1. Alonge, M., Lebeigle, L., Kirsche, M., Jenike, K., Ou, S., Aganezov, S., Wang, X., Lippman, Z.B., Schatz, M.C., Soyk, S.: Automated assembly scaffolding using RagTag elevates a new tomato system for high-throughput genome editing. Genome Biology 23, 258 (2022). 10.1186/s13059-022-02823-7

A2. Cenzato, D., Lipták, Z.: A survey of BWT variants for string collections. Bioinformatics 40(7), btae333 (2024). 10.1093/bioinformatics/btae333

A3. Li, H.: A collection of high-quality human assemblies. Zenodo (2025). 10.5281/zenodo.14854401

A4. Raman, R., Raman, V., Satti, S.R.: Succinct indexable dictionaries with applications to encoding k-ary trees, prefix sums and multisets. ACM Transactions on Algorithms 3(4), 43 (2007). 10.1145/1290672.1290680

